# Biologically-relevant transfer learning improves transcription factor binding prediction

**DOI:** 10.1101/2020.12.21.423873

**Authors:** Gherman Novakovsky, Manu Saraswat, Oriol Fornes, Sara Mostafavi, Wyeth W. Wasserman

**Affiliations:** Centre for Molecular Medicine and Therapeutics, BC Children’s Hospital Research Institute, Vancouver, BC V5Z 4H4, Canada; Department of Medical Genetics, University of British Columbia, Vancouver, BC V6H 3N1, Canada; Department of Statistics, University of British Columbia, Vancouver, BC V6T 1Z4, Canada; Canadian Institute for Advanced Research, CIFAR AI Chair, and Child and Brain Development, Toronto, ON M5G 1M1, Canada

**Keywords:** Transfer learning, deep learning, transcription factor binding prediction, model interpretation

## Abstract

**Background:** Deep learning has proven to be a powerful technique for transcription factor (TF) binding prediction, but requires large training datasets. Transfer learning can reduce the amount of data required for deep learning, while improving overall model performance, compared to training a separate model for each new task.

**Results:** We assess a transfer learning strategy for TF binding prediction consisting of a pre-training step, wherein we train a multi-task model with multiple TFs, and a fine-tuning step, wherein we initialize single-task models for individual TFs with the weights learned by the multi-task model, after which the single-task models are trained at a lower learning rate. We corroborate that transfer learning improves model performance, especially if in the pre-training step the multi-task model is trained with biologically-relevant TFs. We show the effectiveness of transfer learning for TFs with ∼500 ChIP-seq peak regions. Using model interpretation techniques, we demonstrate that the features learned in the pre-training step are refined in the fine-tuning step to resemble the binding motif of the target TF (*i*.*e*. the recipient of transfer learning in the fine-tuning step). Moreover, pre-training with biologically-relevant TFs allows single-task models in the fine-tuning step to learn features other than the motif of the target TF.

**Conclusions:** Our results confirm that transfer learning is a powerful technique for TF binding prediction.

## Background

A subset of human DNA-binding transcription factors (TFs) control gene expression at the transcriptional level by recognizing and binding to specific sequence motifs within *cis*-regulatory regions known as TF binding sites (TFBSs) [1]. The disruption of TF genes and TFBSs is associated with rare genetic disorders [2,3] and cancer [4,5]. Therefore, delineating the regions to which TFs bind in the genome could indicate potential regulatory regions on which to focus analyses, and help to broaden our understanding of how genes are regulated in health and disease.

Chromatin immunoprecipitation followed by sequencing (ChIP-seq) is an experimental assay that enables the identification of TF-bound regions *in vivo* at a resolution of a few hundred base pairs (bp) [6]. These regions, known as ChIP-seq peaks, are expected to be enriched for TFBSs. The ReMap database has compiled and uniformly reprocessed thousands of public ChIP-seq datasets [7,8]. It provides access to millions of ChIP-seq peaks related to the binding of approximately 800 human TFs in 602 different human cell and tissue types. Based on ReMap, the UniBind database stores reliable TFBS predictions from four different computational models, including position weight matrices (PWMs; reviewed in [9]), for the ChIP-seq peaks of 231 human TFs in 315 different human cell and tissue types [10].

Despite large-scale data generation efforts by public consortia such as ENCODE [11], delineating the binding regions of each human TF in the genome remains incomplete. For instance, about 40% of human TFs have not been profiled by ChIP-seq, and only a few, such as CTCF, have been profiled extensively in multiple biological contexts (*e*.*g*. across a range of cell and tissue types). To complement data generation efforts, deep learning methods have become pervasive, as high quality and large-scale datasets have drastically improved their performance (reviewed in [12]). A large training dataset is fundamental to the success of deep learning methods, however, the amount of ChIP-seq data for the majority of human TFs, if available, is small. For example, of the human TFs stored in ReMap, 381 (47.6%) have been profiled in only one cell or tissue type, and 134 (16.7%) have less than 1,000 annotated ChIP-seq peaks.

Transfer learning—reusing the information learned from a model developed for one task as the starting point for a model on a second different, but related, task—has been shown to reduce the amount of data required for training while improving overall model performance for diverse applications (reviewed in [13]). In biology, transfer learning has been successful in several areas, including: reconstructing gene regulatory networks [14–16]; modeling gene expression from single-cell data [17–20]; or predicting genomic features, including accessible regions [21], chromatin interactions [22], and TFBSs [23,24].

The current approach to transfer learning in the field of computer vision consists of two steps: pre-training a model on a large dataset (*e*.*g*. ImageNet [25]), and fine-tuning the model weights to suit a task of interest. The rationale is that in the pre-training step, the model learns low-level image features such as lines or curves [26], which are generalizable to the downstream task. For TF binding prediction, this translates into pre-training a multi-task model with as much genomic data as possible to learn common DNA features (*e*.*g*. TF binding motifs), so that in the fine-tuning step, the downstream task can exploit these common features learned in the pre-training step while focusing on learning novel features specific to that task. It is often unclear what characteristics are responsible for the superior performance of transfer learning. For example, the learning process could reveal common motifs in regulatory regions, simpler DNA sequence composition properties (*e*.*g*. %GC content), or other features, some of which might be novel to our understanding of TF binding specificity. Gaining insights into how the learning process influences predictive performance would allow for a broader adoption and higher impact of transfer learning in genomics.

In this study, we perform an in-depth assessment of transfer learning for TF binding prediction. We corroborate the findings of Zheng and colleagues that transfer learning improves model performance, especially for TFs with small training datasets [24]. We show that transfer learning can perform well even when training with as few as 50 ChIP-seq peaks. We demonstrate that the benefit of transfer learning is greater when pre-training with biologically-relevant TFs. Using model interpretation techniques, we observe that the features learned in the pre-training step are refined in the fine-tuning step to resemble the motif of the target TF (*i*.*e*. the recipient of transfer learning in the fine-tuning step), and pre-training with biologically-relevant TFs allows the model to learn features other than the motif of the target TF in the fine-tuning step, such as the motifs of cofactors. These results advocate for a broader adoption of transfer learning in bioinformatics-related deep learning studies.

## Results

### A sparse matrix of TF binding events across accessible genomic regions

Deep learning-based TF binding prediction can be treated as a binary classification task wherein the ones and zeros (or positives and negatives) represent whether or not a TF binds to a genomic region. It is common to define the regions a TF binds to as the set of ChIP-seq peaks for that TF, with unbound regions being the ChIP-seq peaks from other TFs or randomly selected genomic regions. Leaving aside the intrinsic challenges associated with ChIP-seq data generation and peak-calling [27], there are limitations to adopting such definitions. For instance, many ChIP-seq peaks lack the consensus motif of the profiled TF [28], while others are the consequence of indirect or tethered binding events [29,30], or appear in datasets for unusually high numbers of other TFs [31–33]; *i*.*e*. they may be artefacts. While bound regions can be directly defined from the ChIP-seq data, the selection of unbound regions for use in deep learning models is more difficult. For example, the %GC content, which is often an important contributor to model performance both *in vitro* [34] and *in vivo* [24], varies between the set of peak regions of different TFs [35]. Moreover, some classes of TFs are special: pioneer TFs can bind to nucleosome regions, which have distinctive sequence characteristics [36] and are not typically bound by non-pioneer TFs [37]. Thus, it is suboptimal to define the set of unbound regions for a given TF based on the set of bound regions of other TFs. The alternative approach of using randomly selected genomic regions as unbound regions can result in the inclusion of a set non-relevant regions, such as centromeres or telomeres. Furthermore, a region not bound by a TF in a certain cell or tissue type could be bound by that same TF in a different biological context. Due to the dependence of deep learning on high quality data [38], properties arising from one or more of the aforementioned limitations could result in improperly fitted models, and ultimately, mislead or inflate model performance.

To mitigate these limitations, which in turn could negatively impact on our assessment of transfer learning, we restricted the selection of bound and unbound regions to active regulatory regions of the genome. We constructed a sparse matrix describing the binding of 163 TFs to 1,817,918 200-bp of DNase I hypersensitive sites (DHSs) in a cell and tissue type agnostic manner (**Methods**; **Fig. 1**). Each element in the matrix was defined by a specific TF-DHS pair, and could take one of three values: “1” (*i*.*e*. bound) if the DHS was both accessible and bound by the TF in at least one cell or tissue type in common; “0” (*i*.*e*. not bound) if the DHS was accessible but not bound by the TF in any cell or tissue types in common; or “null” if the binding of the TF to the DHS could not be resolved (*e*.*g*. the TF had not been profiled in a cell or tissue type with available DHS data). The total number of ones, zeros, and nulls in the matrix was ∼1.9M, ∼51.7M, and ∼242.6M, respectively. In addition, the number of ones for each TF varied greatly, with 16 TFs having ≤500 bound regions (**Table S1**).

**Fig. 1:**
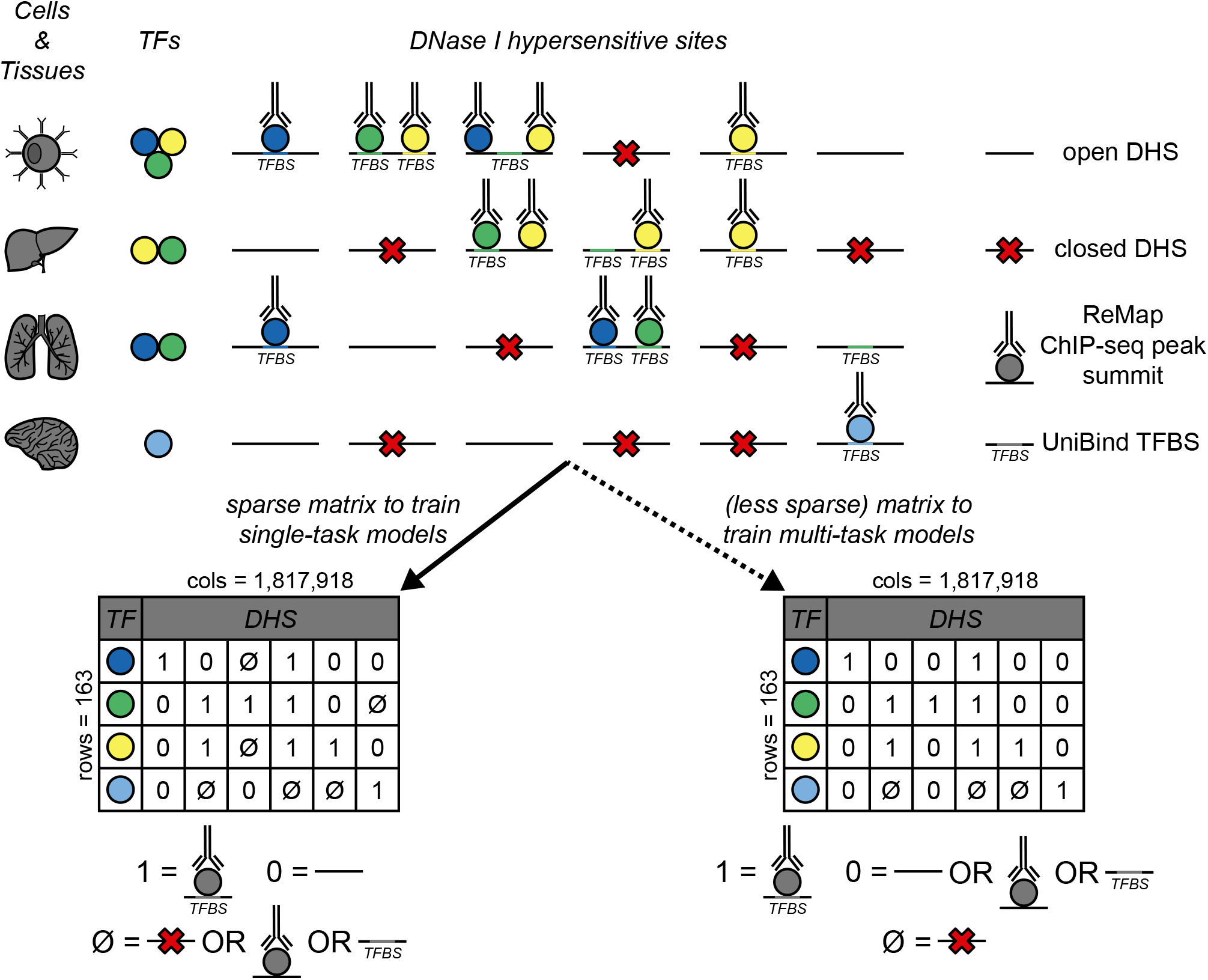
Two matrices of TF binding events across accessible genomic regions. The matrices, one more “sparse” for training single-task models and the other “less sparse” for training multi-task models, summarize the binding (from ReMap ChIP-seq peaks and UniBind TFBSs) of 163 TFs to 1,817,918 DHSs accross 52 cell and tissue types, with rows representing TFs and columns representing DHSs. For each TF-DHS pair in the matrix: a “1” indicates that, according to both ReMap and UniBind, the TF binds to the DHS, and that the DHS is open (*i*.*e*. accessible); a “0”, that the DHS is open but that the TF does not bind to it; and a null sign (“∅”) indicates that there is insufficient evidence to assign a one or a zero, for instance, if the DHS is closed (*i*.*e*. not accessible). Both matrices have the same number of ones, however, they differ in the number of zeros and nulls. In the sparse matrix, TF-DHS pairs wherein the DHS is open and the TF binds to it according to either ReMap or UniBind data (not both) are assigned a null value; instead, in the less sparse matrix, they are assigned a zero value. DHS = DNase I hypersensitive site; TF = transcription factor; TFBS = TF binding site.

The number of unresolved elements (>80%) led to concerns regarding the sparsity of the matrix and whether it had captured known TF-TF functional associations present in ChIP-seq data [30]. We computed pairwise cosine similarities between the binding vectors of all TFs as a measure of correlation (**Methods**). To ease interpretation, TFs were sorted by their hierarchy in TFClass [39] and visualized on a heatmap (**Fig. 2**). As expected, TFs with shared DNA-binding mechanisms (hereafter referred to as binding modes), such as homologs or overlapping members of dimeric complexes, formed well-defined clusters along the heatmap diagonal. In contrast, away from the diagonal of the heatmap, correlated points could indicate the cooperative binding between TFs from different families, such as the erythropoietic TFs GATA1, GATA2, SOX6 and TAL1 [40], highlighted on the heatmap using white arrows. In agreement with previous literature [41], TEAD4 was also highly correlated with these four TFs (teal arrows). To further support the presence of cooperative binding in the TF binding matrix, we compared the binding vector similarities of TF-TF interacting pairs with different degrees of confidence from STRING [42]. On average, the binding vectors of the “highest confidence” TF pairs exhibited a higher degree of similarity than the rest (0.141 vs. 0.063; Welch t-test, *p*-value = 6.46e-09). Taken together, we concluded that the TF binding matrix captured TF cooperativity, and thus was well suited for our assessment of transfer learning.

**Fig. 2:**
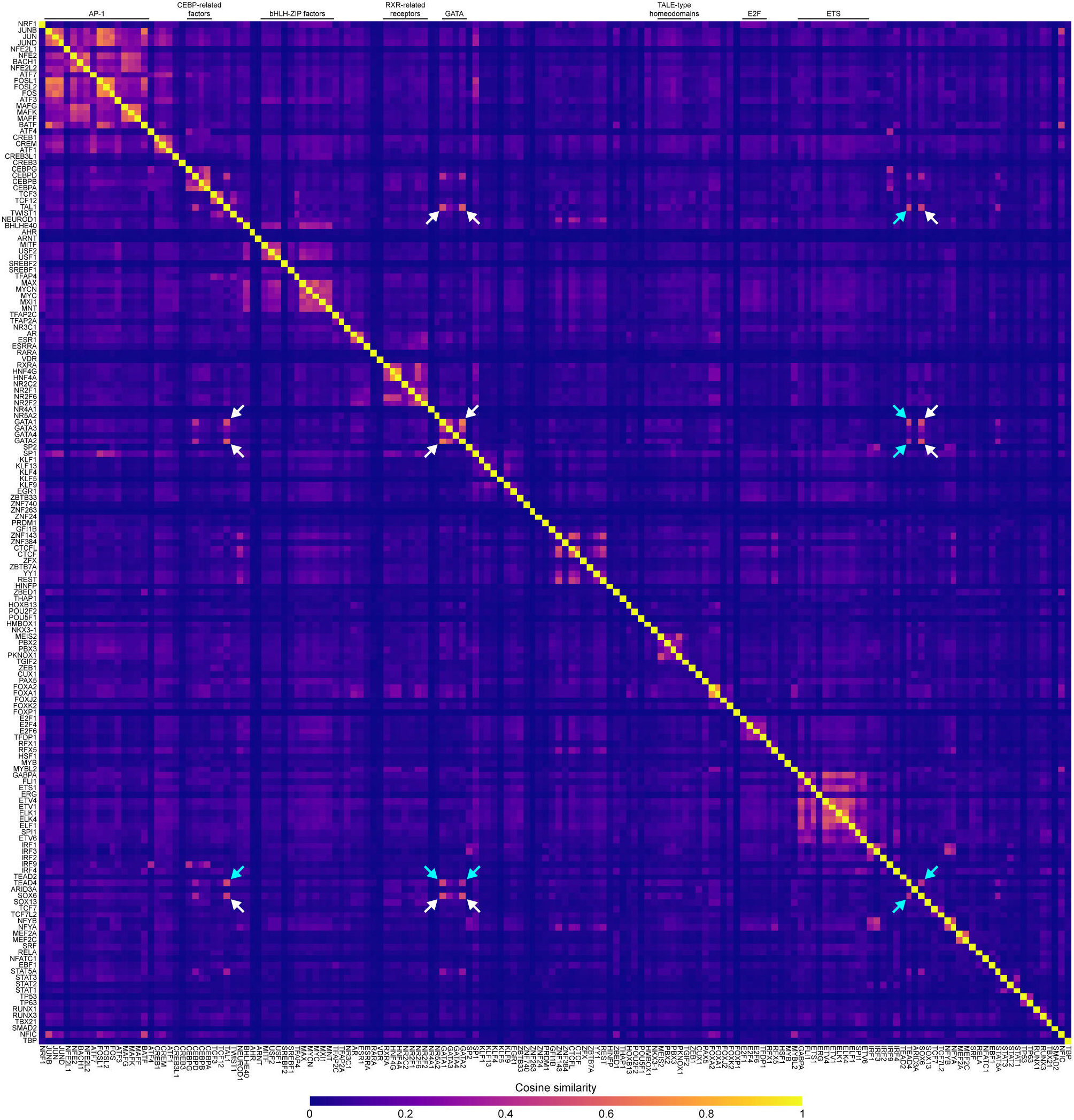
Heatmap of pairwise cosine similarities between the binding vectors of 162 TFs (SMAD3 is excluded because it had no bound regions). Homologous TFs from the same families form well-defined clusters along the heatmap’s diagonal. Highlighted with white arrows, an example of cooperativity for highly correlated TFs from different families, namely GATA1, GATA2, SOX6, and TAL1, which are required for erythropoiesis [40]. TEAD4 is also highly correlated with these four TFs (teal arrows), in accordance with previous literature [41].

### Transfer learning improves TF binding prediction

In a recent preprint, Zheng and colleagues applied transfer learning to predict TFBSs for 38 TFs in GM12878 cells [24]. Their transfer learning framework, named AgentBind, outperformed models trained from scratch (*i*.*e*. without transfer learning), particularly for TFs with little ChIP-seq data. To corroborate these results, we implemented a similar two-step transfer learning strategy for TF binding prediction (**Methods**; **Fig. 3A**). In the pre-training step, a multi-task model (hereafter referred to as multi-model) is trained with multiple TFs; and in the fine-tuning step, single-task models for individual TFs (hereafter referred to as individual models) are initialized with the weights learned by the multi-model, after which they are trained at a lower learning rate. We trained a multi-model using the 50 TFs with the highest number of resolved regions (*i*.*e*. not null) in the TF binding matrix. Moreover, we trained individual models, with and without transfer learning, for 49 of these TFs; ARNT was excluded because it had <250 bound regions left (*i*.*e*. non-overlapping with multi-model regions). Given the advantages of the Matthews correlation coefficient (MCC) over other more popular metrics for evaluating binary classification tasks [43], we chose MCC as our performance metric. Transfer learning significantly improved model performance for 42 (85.7%) TFs (**Fig. 3B**; Wilcoxon signed-rank test, *p*-value = 3.43e-07), especially for TFs with small training datasets, which is in concordance with the results from AgentBind. For the seven cases where transfer learning was detrimental, the differences in model performance compared to training from scratch were small (ΔMCC <0.05; **Table S2**).

**Fig. 3:**
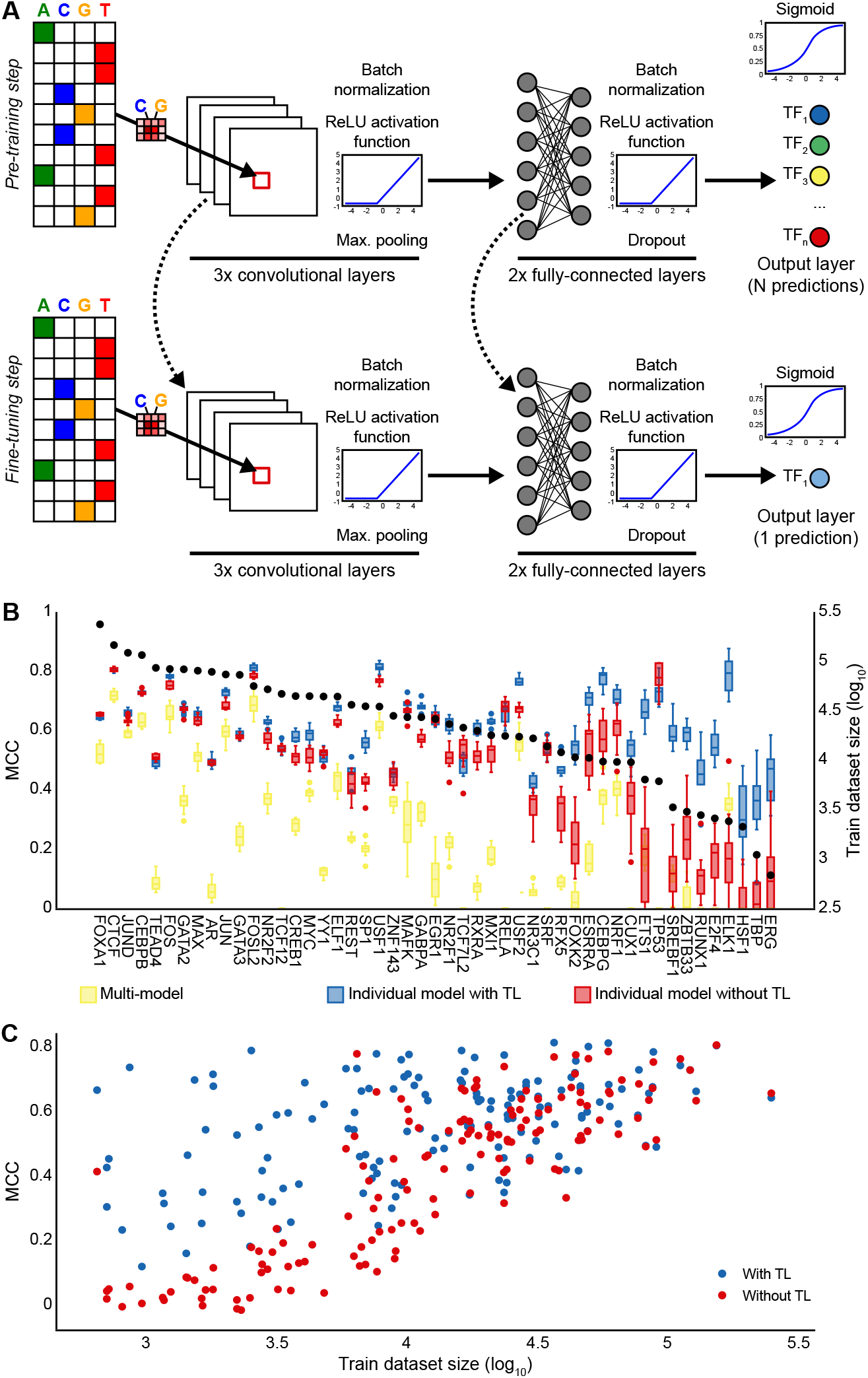
Transfer learning improves TF binding prediction. (**A**) The transfer learning strategy used in this study consists of two steps: pre-training a multi-task CNN with multiple TFs (top); and fine-tuning a single-task CNN, initialized with the weights of the pre-trained multi-task CNN, for one TF (bottom). The CNN architecture is similar to those of Basset [57] and AI-TAC [58]: three convolutional layers, each with ReLU activation, batch normalization, and max-pooling, followed by two fully-connected layers and one output layer. Models were trained with one-hot encoded, 200-bp long DNA sequences and their reverse complements. (**B**) Performance of individual models trained with (blue boxes) and without (red boxes) transfer learning for the 50 TFs from the pre-training step. The performance of each TF on the multi-model is provided as baseline (yellow boxes). (**C**) Quantification of the effect of training dataset size on model performance. TFs (dots) are plotted with respect to the size of their training dataset (x-axis) and performance of their individual models trained with (blue) and without (red) transfer learning (y-axis). CNN = convolutional neural network; MCC = Matthews correlation coefficient; ReLU = rectified linear units; TF = transcription factor; TL = transfer learning.

The pre-training step included data for all TFs, so for a more direct comparison, we repeated the previous experiment with 99 additional TFs that had at least 250 bound regions in the TF binding matrix and, importantly, had not been used to train the multi-model. Again, transfer learning significantly improved model performance and was more beneficial for TFs with small training datasets (**Fig. 3C**; Wilcoxon signed-rank test, *p*-value = 3.98e-13). The most notable example was ATF4, with only 557 bound regions, which achieved a MCC of 0.74 after transfer learning (compared to 0.06 when training without transfer learning). However, transfer learning was detrimental for 17 (17.1%) TFs (**Table S2**). It is worth noting that transfer learning reduced the variation in model performance resulting from different data splits (*i*.*e*. the random assignment of data to different training, validation and test sets), thereby increasing the overall robustness of the training process (**Fig. S1**).

Driven by the observation that transfer learning benefits TFs with small training datasets in particular, we sought to explore the minimum training dataset size required for achieving effective model performance, which we defined as a MCC ≥0.5 (**Methods**). We focused on five TFs from five different families: HNF4A (a nuclear receptor), JUND (a basic leucine zipper), MAX (a basic helix-loop-helix), SPI1 (a tryptophan cluster), and SP1 (a C2H2 zinc finger). For each TF, we trained individual models, with and without transfer learning, by randomly downsampling to 5,000, 1,000, 500, 250, 125, and 50, bound and unbound regions from the TF binding matrix. We repeated the downsampling process five times to ensure the robustness of the results. Models trained with transfer learning achieved effective performance levels when downsampling to just 500 bound/unbound regions (**Fig. 4** for HNF4A; **Figs. S2** to **S5** for the remaining TFs), which was concordant with the previous results for ATF4. For JUND and MAX, transfer learning performance was effective when downsampling to as few as 50 bound/unbound regions (**Figs. S2F** and **S3F**). This could be explained by the use of data for both TFs in the pre-training step. Nevertheless, for SP1, which had also been used to train the multi-model, transfer learning performance fell below effective levels after downsampling to 500 bound/unbound regions (**Fig. S5C**). We attributed this result to the poor performance of SP1 in the multi-model (MCC = 0.20 vs 0.59 and 0.51, for JUND and MAX, respectively; **Table S2**), suggesting that the multi-model could not correctly learn the binding features of SP1, which in turn would result in an inadequate set of weights with which to initialize the individual model of SP1 in the fine-tuning step.

**Fig. 4:**
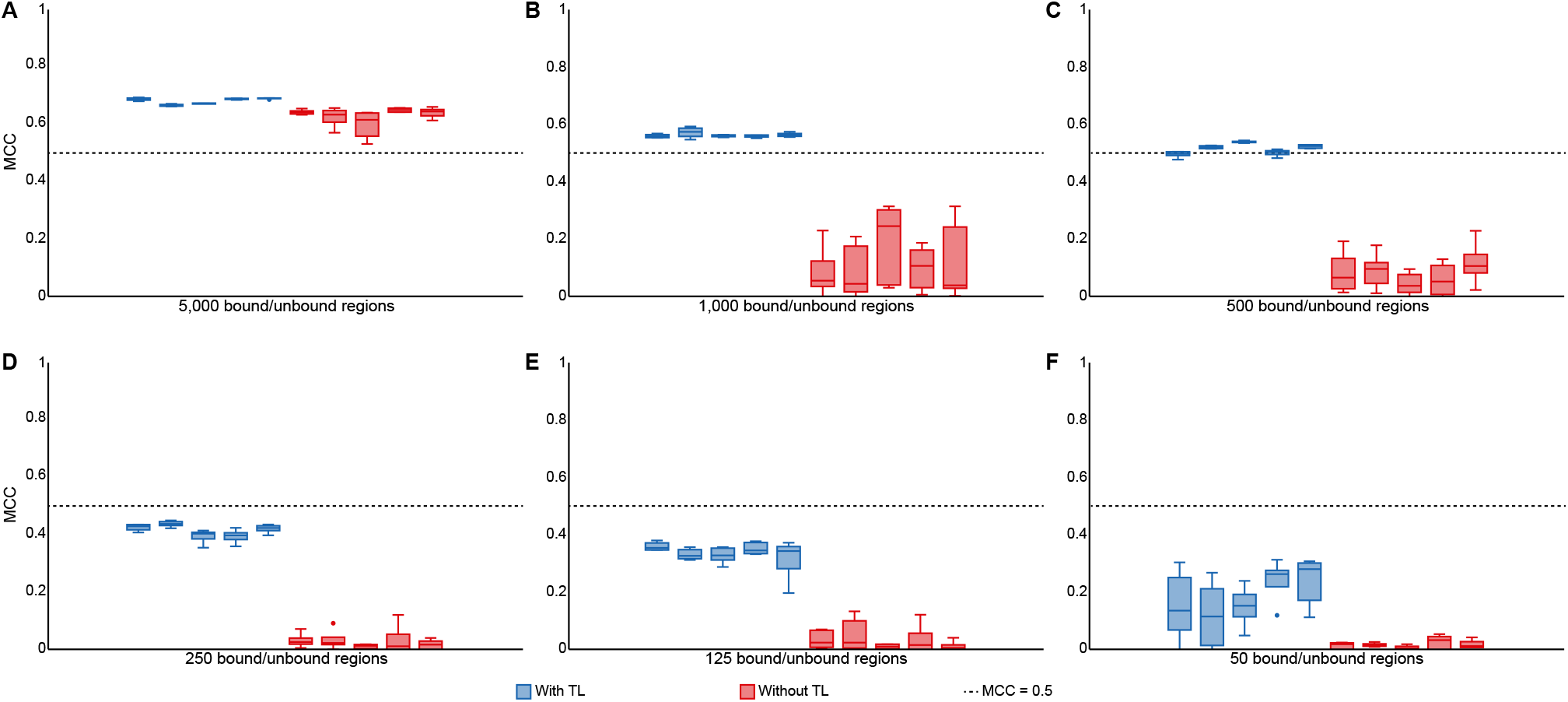
Transfer learning performs well for TFs with small training datasets. Performance of HNF4A models trained with (blue boxes) and without (red boxes) transfer learning on 5,000 (**A**), 1,000 (**B**), 500 (**C**), 250 (**D**), 125 (**E**), and 50 (**F**), bound and unbound regions. Each model was trained five times with different random initializations to ensure the robustness of the results. MCC = Matthews correlation coefficient; TF = transcription factor; TL = transfer learning.

### Biologically-relevant prior knowledge improves transfer learning

TFs from the same family often have highly similar DNA-binding specificities [44]; hence, we hypothesized that the presence in the pre-training step of TFs with the same binding mode as the target TF could have a positive effect on transfer learning performance. The JASPAR database of TF binding profiles includes a hierarchical clustering that groups TFs based on the similarity of their binding profiles [45]. We relied on this grouping as the source of binding modes (**Table S3**). Focusing on the set of TFs whose performance worsened with transfer learning, we found that the binding modes of the most extreme cases (ΔMCC > 0.05), namely MEF2A, HOXB13, MEF2C, TFAP2A, and ZNF384, differed from those of the 50 TFs used to train the multi-model, suggesting that the inclusion in the pre-training step of TFs with relevant binding modes could be beneficial. This observation was further supported by reviewing the remaining TFs; TFs with the same binding mode as a TF from the pre-training step benefited from transfer learning significantly more (Welch t-test, *p*-value = 0.01; **Fig. 5**).

**Fig. 5:**
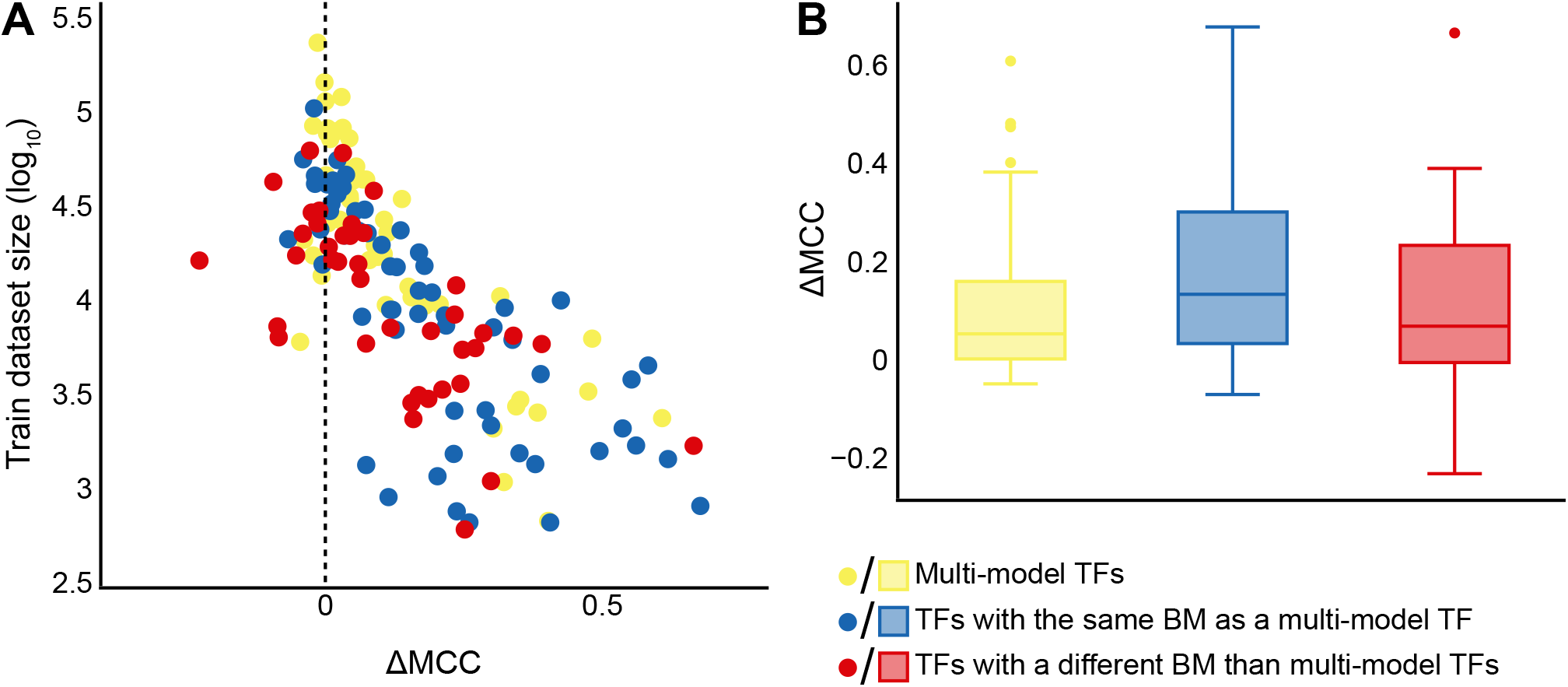
Transfer learning is more beneficial for TFs with the same binding mode as a TF from the pre-training step. (**A**) Performance difference (*i*.*e*. ΔMCC) of individual models trained with and without transfer learning (x-axis) is plotted with respect to the size of the training dataset (y-axis) for TFs used to train the multi-model (yellow dots), TFs with the same binding mode as a multi-model TF (blue dots), and TFs with different binding modes than multi-model TFs (red dots). (**B**) ΔMCCs for TFs used to train the multi-model (yellow boxes), TFs with the same binding mode as a multi-model TF (blue boxes), and TFs with different binding modes than multimodel TFs (red boxes). BM = binding mode; MCC = Matthews correlation coefficient; ΔMCC = MCC from transfer learning - MCC from training from scratch; TF = transcription factor.

We hypothesized that the presence in the pre-training step of other biologically-relevant prior knowledge, such as cofactors, or functional partners from STRING, could also have a positive effect on transfer learning performance (**Methods**). We defined cofactors as pairs of TFs whose binding was positively correlated. We focused on the same pentad of TFs from the previous section (*i*.*e*. HNFA4, JUND, MAX, SPI1, and SP1). For each TF, we pre-trained five different multi-models with: five TFs with the same binding mode as the target TF; five cofactors of the target TF; five TFs with the same binding mode as the target TF but whose binding was not correlated with it (*i*.*e*. non-cofactors); five functional partners of the target TF from STRING; and five randomly selected TFs. Cofactors, functional partners from STRING, and randomly selected TFs were restricted to have different binding modes than the target TF. To avoid any confounding effects related to the training dataset size, all models were trained with a similar number of regions: ∼70,000 for multi-models and ∼2,000 for individual models. Furthermore, to set an upper performance limit for each pre-training strategy, we repeated the analysis by replacing one of the five TFs, with which we pre-trained the multi-model, by the target TF. As expected, pre-training with the target TF resulted in better transfer learning. Moreover, when the target TF was included in the multi-model, we did not observe any significant differences between the five pre-training strategies (Kruskal-Wallis H-test, Bonferroni adjusted *p*-values = NS; **Fig. 6**). In contrast, when the target TF was not included in the multi-model, pre-training with either TFs with the same binding mode as the target TF or with cofactors were the best strategies: both achieved effective performance levels for four out of five TFs (except for SP1). Interestingly, using non-cofactors with the same binding mode as the target TF during pre-training led to slightly worse performance, suggesting that cofactors could play an important role in TF binding prediction. Of the five TFs analyzed, SP1 was a notable outlier: not only did it show the worst performance levels overall, but when it was not included in the multi-model, pre-training with other Krüppel-like zinc fingers sharing the same binding mode as SP1 performed worse than pre-training with randomly selected TFs (**Fig. 6E**). To confirm whether these observations were general to the Krüppel-like zinc finger family of TFs, we repeated the experiment with another Krüppel-like zinc finger, EGR1. We obtained results more in line with the other four TFs (**Fig. 6F**), suggesting that SP1 was an isolated case. Finally, to ensure that these results were not biased by our choice of model architecture, we repeated the previous experiment using the hybrid architecture with convolutional and recurrent layers of DanQ [46], obtaining similar results (**Fig. S6**). Consistent with previous studies [24,46], model performance was slightly better when using the DanQ architecture.

**Fig. 6:**
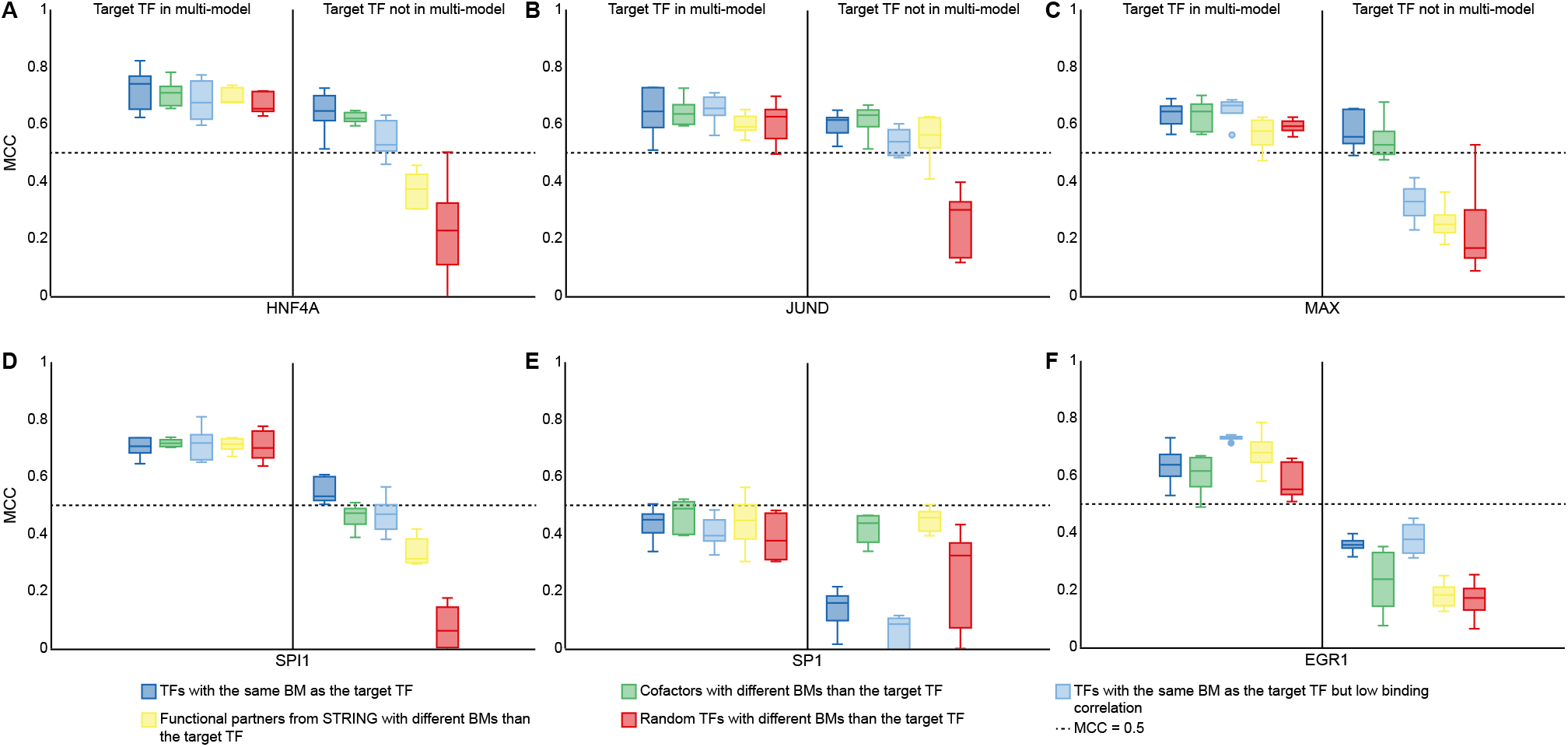
Biologically-relevant transfer learning improves model performance. Transfer learning performance for the target TFs HNF4A (**A**), JUND (**B**), MAX (**C**), SPI1 (**D**), SP1 (**E**), and EGR1 (**F**), from multi-models pre-trained with five TFs with the same binding mode as the target TF (dark blue boxes), five cofactors of the target TF with a different binding mode than the target TF (green boxes), five non-cofactors with the same binding mode as the target TF (light blue boxes), five functional partners of the target TF from STRING with a different binding mode than the target TF (yellow boxes), and five randomly selected TFs with a different binding mode than the target TF (red boxes), with (left) and without (right) the presence of the target TF in the pre-training step. BM = binding mode; MCC = Matthews correlation coefficient; TF = transcription factor.

The pentad TFs had a sufficiently large number of bound regions that we downsampled; hence, we wondered whether TFs with less bound regions would behave similarly. For each of the pentad TFs, we selected two additional TFs with the same binding mode and within the following ranges of bound regions: <1,000 positives or 1,000-10,000 positives. For each of these 15 TFs (the five “pentad” TFs plus the 10 newly selected), we trained individual models using transfer learning from two different multi-models pre-trained with either five TFs with the same binding mode as the target TF, or five randomly selected TFs with different binding modes than the target TF. Each multi-model was trained both with and without the target TF included in the pre-training step. Unlike the above set of experiments, this time we did not perform downsampling. In general, pre-training with the target TF resulted in better transfer learning performance (**Fig. 7**), however, for TFs with large datasets, this effect was less obvious. In addition, pre-training with a biologically-relevant multi-model improved transfer learning performance, even when the target TF was present in the multi-model, especially for TFs with <10,000 positives. Similar results were obtained when the experiment was repeated, but pre-training using five cofactors (**Fig. S7**).

**Fig. 7:**
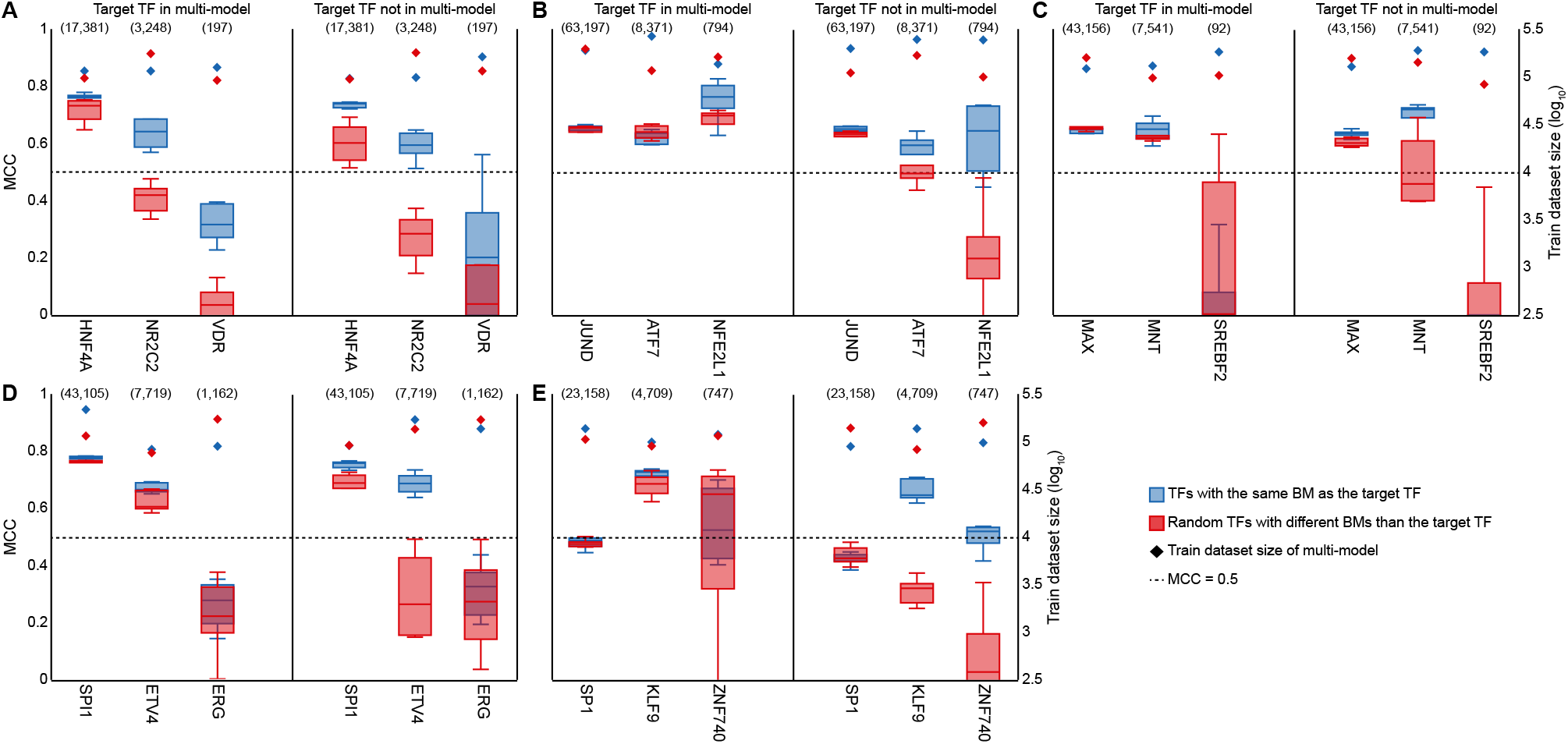
The use of relevant binding mode information in the pre-training step improves model performance for TFs with small datasets, regardless of their presence in the multi-model. Transfer learning performance for three target TFs with the same binding mode, but different number of bound regions, from each of the following five families: (**A**) HNF4A, NR2C2, and VD2R (nuclear receptors); (**B**) JUND, ATF7, and NFE2L1 (basic leucine zippers); (**C**) MAX, MNT, and SREBF2 (basic helix-loop-helix factors); (**D**) SPI1, ETV4, and ERG (tryptophan cluster factors); and (**E**) SP1, KLF9, and ZNF740 (C2H2 zinc fingers). Transfer learning performance for each target TF from multi-models pre-trained with five TFs with the same binding mode as the target TF (dark blue boxes), and five randomly selected TFs with a different binding mode than the target TF (red boxes), with (left) and without (right) the presence of the target TF in the pre-training step. The training dataset size of each multi-model is indicated with diamonds (secondary y-axis). The number of bound regions for each TF is shown between parenthesis. BM = binding mode; MCC = Matthews correlation coefficient; TF = transcription factor.

Next, we tested whether focusing on a few biologically-relevant TFs, rather than pre-training with a large dataset, would be a more effective transfer learning strategy. For each category of biologically-relevant information (*i*.*e*. binding modes, cofactors and functional partners from STRING), we pre-trained two different multi-models for each of the pentad TFs with either: five biologically-relevant TFs; or “burying” the same five TFs amongst 45 TFs drawn from the original multi-model, each with a different binding mode than the target TF. For this set of experiments, the target TF was not included in the pre-training step, and we fixed the training datasets sizes of multi-models and individual models to ∼40,000 and ∼2,000 regions, respectively. Furthermore, we used cofactors and functional partners from STRING with and without sharing the same binding mode as the target TF. We observed that pre-training with just five biologically-relevant TFs led to only slightly better performance levels (**Fig. 8**).

**Fig. 8:**
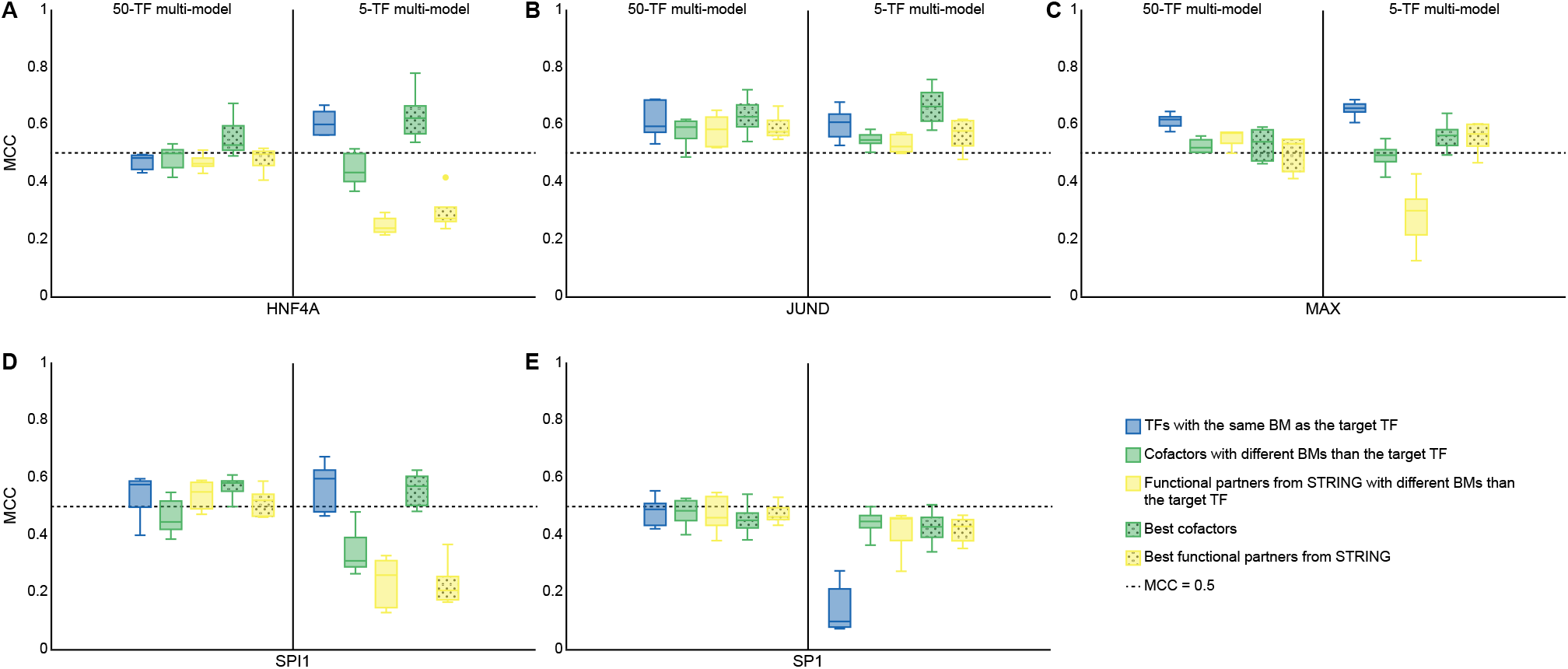
Pre-training with a few biologically-relevant TFs only offers a slight advantage over a multi-model pre-trained with a large number of TFs. Transfer learning performance for the target TFs HNF4A (**A**), JUND (**B**), MAX (**C**), SPI1 (**D**), and SP1 (**E**): on the right, from multi-models pre-trained with five TFs with the same binding mode as the target TF (dark blue boxes), five cofactors of the target TF with a different binding mode than the target TF (green boxes), five functional partners of the target TF from STRING with a different binding mode than the target TF (yellow boxes), and the best cofactors and functional partners from STRING of the target TF regardless of their binding mode (dotted green and yellow boxes, respectively); on the left, from “burying” each group of five biologically-relevant TFs amongst 45 TFs drawn from the original multi-model, each with a different binding mode than the target TF. BM = binding mode; MCC = Matthews correlation coefficient; TF = transcription factor.

### Interpretation of the transfer learning mechanism

In an attempt to understand the mechanism of transfer learning, we converted the filters from the first convolutional layer of the original multi-model to PWMs, and compared them using Tomtom [47] to TF binding profiles from JASPAR (**Methods; Fig. 9A**). As expected, the majority of filters (55%) had significant similarities to known JASPAR motifs (Tomtom *q*-value <0.05; **Table S4**). Focusing on HNF4A, which had not been used in the pre-training step, we found that four of the multi-model filters were refined in the fine-tuning step to resemble the motifs of HNF4A (**Fig. 9B**). We also observed that an increased number (six) of filters became similar to the motifs of HNF4A after using transfer learning compared to training from scratch (three; **Table S4**). Moreover, using transfer learning, the individual model of HNF4A was able to learn the two distinct binding modes of HNF4A represented by the JASPAR profiles MA0114.4 and MA1494.1. Similar observations were made for SPI1, which was also not present in the pre-training step (**Fig. 9B**; **Table S4**). Taken together, these findings suggest that weights from the pre-training step provide a better initialization for convolutional filters in the fine-tuning step, which are then refined to resemble the binding motif of the target TF. This would, in turn, explain the increased performance of transfer learning compared to training from scratch. To confirm that the refinement of convolutional filters in the fine-tuning step was responsible for the improvement in model performance by transfer learning, we applied an alternative fine-tuning strategy: we froze the convolutional layers of the pre-trained multi-model and trained only on the fully-connected layers (*i*.*e*. no refinement of convolutional layer filters was allowed). Doing so resulted in poorer model performance, particularly for TFs that were not present in the multi-model (**Table S5**), further supporting the importance of filter refinement in the fine-tuning step.

**Fig. 9:**
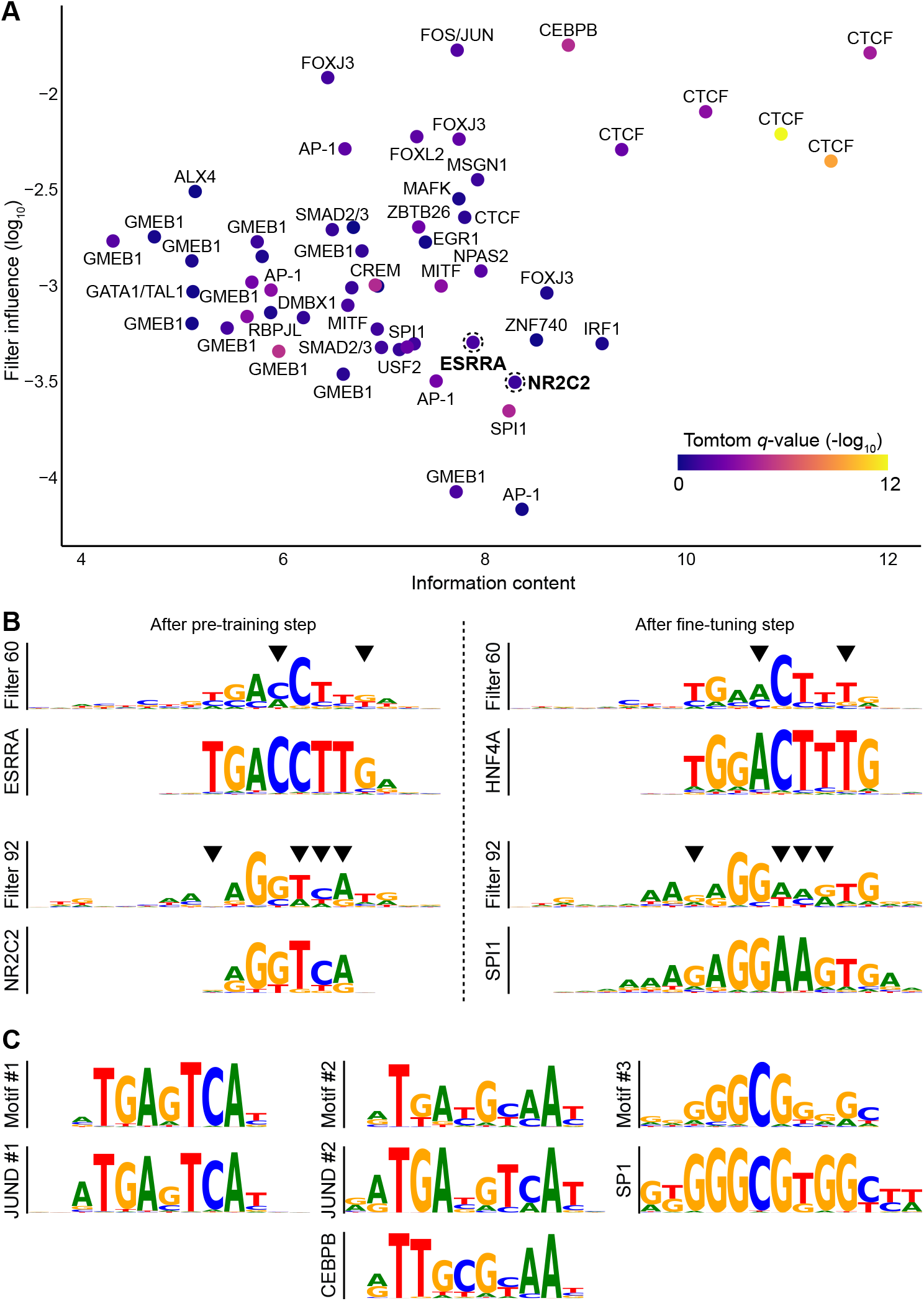
Model interpretation elucidates the transfer learning mechanism. (**A**) TF motif representations learnt in the first convolutional layer of the original multi-model of 50 TFs. Filters (dots) are plotted with respect to their information content (x-axis) and overall influence (y-axis). The color-scale reflects the Tomtom similarity of a given filter to a TF profile from the JASPAR database. When applicable, the name of the most significant TF is shown. Filters 60 and 92 correspond to the JASPAR profiles for ESRRA and NR2C2 (shown in bold), respectively, and are highlighted with dotted line circles. (**B**) Transfer learning fine-tunes the filters learned in the pre-training step to resemble the binding motif of the target TF. Specific positions (black triangles) from filters 60 and 92 (highlighted above) are refined in the fine-tuning step to resemble the JASPAR profiles for HNF4A and SPI1, respectively. (**C**) Learning the motifs of cofactors contributes to the predictive capacity of the model. Using DeepLIFT importance scores from a JUND model initialized with the weights of a multi-model pre-trained with five cofactors, TF-MoDISco recovers the motifs of two of the cofactors, CEBPB and SP1, as well as the JUND motif.

Finally, we applied DeepLIFT [48] and TF-MoDISco [49] in an attempt to understand the role of cofactors in transfer learning. Briefly, DeepLIFT quantifies the importance to the model prediction of each nucleotide in a given genomic sequence, and TF-MoDISco clusters “important” nucleotides from different genomic sequences into motifs. We show an analysis of transfer learning for JUND from two different multi-models pre-trained on either: five cofactors with a different binding mode than JUND (CEBPB, MAFF, MAFG, NFIC and SP1); or five TFs from five different families whose binding was not correlated with JUND (CTCF, EBF1, MXI1, NFYA and TCF3). Data for JUND was not present in the pre-training step. The three motifs generated by TF-MoDISco for the cofactor model corresponded to the canonical JASPAR motifs of CEBPB and SP1, both of which were amongst the cofactors used to train the multi-model, as well as JUND (**Fig. 9C**). For the second model trained on five unrelated TFs, TF-MoDISco identified five motifs, none of which corresponded to the canonical JUND motif. Taken together, this suggests that the model uses the presence of CEBPB and SP1 motifs to aid in the prediction of JUND binding.

## Discussion

In this study, we have demonstrated how incorporating prior knowledge via transfer learning can improve TF binding prediction in deep learning methods, with notable benefits for TFs with scarce experimental binding data. We constructed a large data matrix of TF binding events through the combination of DNA accessibility with experimental and computational TF binding information to define TF-bound and unbound regions with high confidence. Using this matrix, we implemented a two-step transfer learning approach for TF binding prediction that first draws from data from a large number of TFs to learn the general properties of regulatory regions, and second, exploits these properties to generate a specific deep learning model for a single TF. We introduced methods that allowed us to study the learned properties in the deep learning models, providing insight into the transfer learning mechanism. We confirm not only the benefits of including binding data from homologs of the target TF into the first step of transfer learning (consistent with studies predating deep learning [50]), but also the importance of including cofactors. Finally, when focussing specifically on TFs for which experimental binding data is scarce, we show that transfer learning is routinely successful in generating a deep learning model from only 500 experimental regions, and in a few extreme cases it can be successful from as few as 50 regions.

While other studies have reported similar findings about the benefits of transfer learning for TF binding prediction [23,24], we demonstrate that the effectiveness of transfer learning depends largely on the functional association between the target TF and those included in the pre-training step. We show improvements when pre-training with TFs that have the same binding mode as the target TF compared to pre-training with randomly selected TFs. Similar benefits are observed for other biological categories such as cofactors or, to a lesser extent, functional partners from STRING. However, for TFs with large datasets, if the target TF is included in the multi-model, then transfer learning performance is similar for biologically-relevant and randomly selected pre-training sets of TFs. For TFs with small training datasets, the opposite occurs, even when the target TF is included in the multi-model. One possible explanation is that if a TF has enough training examples, then the model can infer its binding without any additional aid (from cofactors or binding mode information). In contrast, when the target TF has few bound regions (relative to other TFs), their inclusion in the multi-model stage will result in the inability of the model to learn relevant features of the target TF for subsequent fine-tuning. In this situation, the inclusion of biologically-relevant information (*e*.*g*. cofactors) may be of benefit.

The idea of using biologically-relevant prior knowledge of the target TF in the pre-training step is enticing, however, it can be challenging to do so. Training a separate multi-model for each TF is computationally expensive. Furthermore, obtaining prior knowledge for a specific TF could prove difficult. For instance, it may not be straightforward to identify cofactors for a TF with few ChIP-seq peaks, as its binding vector may not correlate well with those of other TFs. Binding mode information may be unavailable for new TFs or for TFs from families with fewer members. Lastly, the STRING data available for some TFs could be of low confidence. With these limitations, one might consider pre-training a larger multi-model with most, if not all, of the existing binding modes. Such a multi-model would be pre-trained with the best representative TF (*i*.*e*. the TF with the largest training dataset) from each binding mode, would be more generalizable to other TFs, and would avoid having to pre-train a separate multi-model for each specific case. While our findings, particularly those from the experiment wherein we “buried” five biologically-relevant TFs in 45 additional TFs, show that in most cases pre-training on a few biologically-relevant TFs results in better transfer learning, the generalized approach with 50 TFs contributing to the multi-model performed nearly as well. Optimizing the size and properties of the pre-training dataset to maximize binding mode diversity is likely to be a fruitful avenue of future research.

For image recognition, deep learning models tend to learn simple and basic features in the first layers (lines, curves, etc.), whereas more complex and resolved features emerge at deeper layers. Therefore, it is commonplace to train new models by taking big pre-trained models (multi-models in our case) and freezing the initial layers, focusing the fine-tuning onto the deeper layers. We demonstrate that such an approach doesn’t work for TF binding, as it is likely that the main features, such as TF binding motifs, are already learned in the first convolutional layers.

We have not explored the impact of the learning rate on transfer learning performance. Often, a small learning rate is applied when one uses a pre-trained model based on a CNN. However, an extremely low learning rate makes it challenging for the model to learn new features. In contrast, with high learning rates, weights from the pre-trained model can be ignored, resulting in the loss of the prior knowledge. In this study, when fine-tuning individual models, we used a learning rate 10 times lower than that used to train multi-models. Further exploration of the impact of learning rate on resolving motifs from the data may be a focus of future studies. As for the learning rate, the batch size parameter should also be explored as recent studies have highlighted its impact on model performance [51] and transfer learning efficacy [52].

In conclusion, transfer learning is a powerful technique for TF binding prediction, and we anticipate that it will become essential for the study of TF binding and in other areas of biology.

## Conclusions

Existing deep learning frameworks readily allow the transfer of learned properties between models within the same architecture, overcoming the computational cost and reducing the amount of training data required. This study is a demonstration of one approach to improving the performance of deep learning models for TF binding prediction via transfer learning through the incorporation of field-/domain-specific information. Our results advocate for a broader adoption of such focused transfer learning strategies in deep learning, particularly for biological sequence analysis.

## Methods

### TF binding matrices

We built two TF binding matrices (*i*.*e*. data structures containing information about TF binding events, not motif models), one more sparse for training individual models and the other (less sparse) suitable for training multi-models. The matrices aggregate the binding data of 163 TFs for 1,817,918 accessible genomic regions accross 52 cell and tissue types, with rows representing TFs and columns representing accessible genomic regions (**Fig. 1**). As the source of TF binding events, we used ChIP-seq peak summits from ReMap 2018 [7] and PWM-based TFBS predictions from UniBind [10]. We used the track of ENCODE DHS peak clusters from the UCSC Genome Browser [53,54] as the accessible genomic regions. Regions were resized to 200 bp around the center of each DHS cluster using bedtools slop [55]. Data was matched by cell and tissue type. For the sparse matrix (*i*.*e*. for training individual models), each element (*i*.*e*. a TF-DHS pair) was assigned one of three values: “1” (*i*.*e*. bound) if the region was accessible (*i*.*e*. DHS positive) and overlapped with both ReMap (*i*.*e*. ChIP-seq peak positive) and UniBind (*i*.*e*. TFBS positive) in at least one matched cell or tissue type; “0” (*i*.*e*. unbound) if the region was accessible but did not overlap with both binding features of the TF in any matched cell and tissue type; or “null” if the binding of the TF to the region could not be resolved. A null value indicated insufficient evidence to determine if the region was or was not bound by the TF: the region may not be accessible (*i*.*e*. DHS negative) in any matched cell or tissue type for which the TF binding had been profiled, or it may be accessible but only overlap with one type of binding feature of the TF (i.e. ReMap-peak or UniBind motif, but not both). For the less sparse matrix (*i*.*e*. for training multi-models), unresolved elements (*i*.*e*. null) wherein the region was accessible but only overlapped with one type of binding feature of the TF (not both), were instead assigned a value of 0 (*i*.*e*. unbound) to make it suitable for size reduction when training multi-models.

### Cosine similarity

The binding vector of a TF was given by the row in the sparse matrix corresponding to that TF. For a pair of TFs, the cosine similarity between their binding vectors was computed using scikit-learn [56]. Unresolved regions of either TF were removed from both vectors prior to calculation.

### Model architecture

We adapted the CNN architecture from Basset [57] and AI-TAC [58] for TF binding prediction:

- 1st convolutional layer with 100 filters (19×4), batch normalization, ReLU activation, 0% dropout and max pooling (3×3);
- 2nd convolutional layer with 200 filters (7×1), batch normalization, ReLU activation, 0% dropout and max pooling (3×3);
- 3rd convolutional layer with 200 filters (4×1), batch normalization, ReLU activation, 0% dropout, and max pooling (3×3);
- 1st fully connected layer with 1,000 nodes, batch normalization, ReLU activation, and 30% dropout;
- 2nd fully connected layer with 1,000 nodes, batch normalization, ReLU activation, and 30% dropout; and
- Fully-connected output layer with 1, 5 or 50 outputs (depending on the model).

For DanQ, we used the following specifications:

- 1st convolutional layer with 320 filters (26×4), ReLU activation, 20% dropout and max pooling (13×13);
- 2 bi-directional LSTM layers with hidden state size 320 and 50% dropout;
- 1st fully-connected layer with 925 nodes and ReLU activation; and
- Fully-connected output layer with 1, 5 or 50 outputs (depending on the model).

Both architectures were implemented using the PyTorch framework [59].

### Transfer learning

We implemented a two-step transfer learning strategy similar to the one used in AgentBind [24]. In the pre-training step, we trained a multi-model. In the fine-tuning step, we initialized the model of the target TF by transferring all of the layers learned by the multi-model, except the output layer. We then trained the initialized model of the target TF with a 10-fold lower learning rate (*i*.*e*. 0.0003). In the fine-tuning step, we either train the entire model, or just the fully-connected layers.

To train multi-models, we used the less sparse matrix. We extracted a slice of the matrix containing the row vectors of all TFs included in the multi-model. Any column vectors containing unresolved elements were removed. Then, we randomly split the regions into training (80%), validation (10%), and test (10%) sets using scikit-learn. The training and validation sets additionally included the reverse-complement of each region. We trained the model using the Adam optimizer [60]. We applied one-hot encoding to convert nucleotides into 4-element vectors (sequences with Ns were discarded), set the learning rate to 0.003 and batch size to 100, and used an early stopping criteria to avoid overfitting when the model performance on the validation set did not improve.

Individual models were trained in a similar way, but using the sparse matrix. The number of bound versus unbound regions for all TFs was imbalanced. For example, one of the most abundant TFs in the matrix, CTCF, had 1,656,242 resolved regions of which <5% were bound. To deal with the imbalance, we downsampled the set of unbound regions to a 50:50 ratio while accounting for the %GC content distributions between bound and unbound regions. To ensure the robustness of our results, each individual model was trained five times with different, randomly generated training/validation/test splits. To avoid overfitting, there was no overlap between the regions used in the pre-training and fine-tuning steps within the same experiment.

### Model performance

Model performance was evaluated using the MCC metric, calculated with scikit-learn. We compared the area under the receiver operating characteristic (AUCROC) and precision-recall (AUCPR) curves to the MCC metric obtained for 148 individual models trained with transfer learning (**Fig. S8**). We defined effective model performance by AUCROC and AUCPR of ∼0.835, which corresponded to MCC >= 0.5. For reference, when trained on 200-bp long sequences and evaluated on chromosomes 8 and 9, DeepSEA was reported to achieve an average AUCROC of ∼0.87 [61].

### Pre-training with biologically-relevant TFs

The pentad TFs used in the analyses (*i*.*e*. HNFA4, JUND, MAX, SP1, and SPI1) belonged to the following binding modes: 2 and 4 (HNF4A); 1 and 18 (JUND); 7 (MAX); 34 (SP1); and 16 (SPI1). At least five additional TFs in the sparse matrix shared one or more binding modes with a pentad TF. For multi-models trained with the target TF, we randomly selected four other TFs sharing one or more binding modes with the target TF (five when training without the target TF).

To identify potential cofactors based on correlated binding, we computed pairwise cosine similarities between the binding vectors of 162 TFs (SMAD3 was removed because it did not have any bound regions in the sparse matrix). For a given TF, we sorted the remaining 161 TFs by cosine similarity, and removed those sharing one or more binding modes with it. Out of the remaining TFs, we selected the top five for multi-model training. If the multi-model was trained with the target TF, we only selected the top four TFs. When focusing on the best cofactors in **Figs. 7** and **S7**, we kept all TFs after sorting. Similarly, to select TFs with low binding correlation that shared one or more binding modes with the target TF, we removed those with different binding modes than the target TF after sorting the TFs by cosine similarity, and selected the bottom five TFs. Again, when training the multi-model with the target TF, only the bottom four TFs were selected.

The STRING database stores known and predicted protein-protein interactions, and provides a confidence score for the interactions; we used version 11.0 [42]. We defined the functional partners of a TF as its set of interactors from STRING. To pre-train with prior knowledge from STRING-based associations, we sorted the functional partners of the target TF by confidence score, and removed those that shared one or more binding modes with it. The top five TFs were selected for pre-training (the top four if pre-training with the target TF). As with selecting the best cofactors, when focusing on the best functional partners from STRING in **Figs. 7** and **S7**, we kept all TFs.

Finally, to pre-train on random TFs, we randomly selected five TFs (or four if pre-training with the target TF) with different binding modes than the target TF.

### Model interpretation

To interpret the performance of the original multi-model, we converted each of the 100 filters from the first convolutional layer to PWMs, as in AI-TAC. For each filter, we constructed a position frequency matrix (PFM) from all 19-mers (*i*.*e*. DNA sequences of length 19) that activated that filter by ≥50% of its maximum activation value in all correctly predicted regions. PFMs were then converted to PWMs using scripts from [58], and the background uniform nucleotide frequency was set to 0.25. The resulting PWMs were compared to vertebrate profiles from the JASPAR 2020 database using Tomtom (version 5.0.5) [47].

The influence of each filter in the multi-model was obtained by “silencing” that filter and computing the impact on the model’s predictive capacity. Silencing was achieved by setting the activation values of the filter across all samples in the batch to zero. The resulting output was passed through the remaining layers of the model to obtain the prediction values after silencing. Using this approach, we computed the average influence value for each filter in the model by averaging the square of the differences between the actual and silenced predictions.

We generated DeepLIFT [48] importance scores with ten reference sequences for each positively predicted sample in the test set using the Captum library [62]. To obtain motifs from DeepLIFT importance scores, we used TF-MoDISco [49] with default settings.

## Abbreviations

AUCPR: area under the precision-recall curve
AUCROC: area under the receiver operating characteristic curve
BM: binding mode
ChIP-seq: chromatin immunoprecipitation followed by sequencing
CNN: convolutional neural network
DHS: DNase I hypersensitive site
MCC: Matthews correlation coefficient
PFM: position frequency matrix
PWM: position weight matrix
TL: transfer learning
TF: transcription factor
TFBS: transcription factor binding site

## Declarations

### Ethics approval and consent to participate

Not applicable.

### Consent for publication

Not applicable.

### Availability of data and materials

The TF binding matrices, together with the source code for generating them, are available as 2D numpy arrays [63] on GitHub at https://github.com/wassermanlab/TF-Binding-Matrix. The IPython notebooks and scripts for performing the different transfer learning experiments are also available on GitHub at https://github.com/wassermanlab/TF-Binding-Transfer-Learning.

### Author’s contributions

G.N., M.S., and O.F. devised the project with contributions from S.M. and W.W.W.; G.N. and M.S. performed all experiments; O.F. created the data matrices and performed all statistical analyses; G.N., M.S., O.F., and W.W.W. wrote the manuscript.

## Acknowledgements

We thank the members of the Mostafavi and Wasserman labs for proof-reading the manuscript and providing useful feedback, in particular Alice M. Kaye, Alexandra (Sasha) Maslova, Etienne Meunier, Chendi Wang, and Solenne Correard. We thank Dora Pak and Jonathan Chang for administrative and IT support, respectively.

## Authors’ information

S.M. present address: Paul G. Allen School of Computer Science & Engineering at the University of Washington, Seattle, WA 98195-2350, USA.

## Figures

**Fig. S1**: Performance variance (*i*.*e*. σ; y-axis) of individual models trained with (blue) and without transfer learning (red) is plotted with respect to the size of the training dataset (x-axis) for 148 TFs. MCC = Matthews correlation coefficient; TF = transcription factor; TL = transfer learning.

**Fig. S2**: Performance of JUND models trained with (blue boxes) and without (red boxes) transfer learning on 5,000 (**A**), 1,000 (**B**), 500 (**C**), 250 (**D**), 125 (**E**), and 50 (**F**), bound and unbound regions. Each model was trained five times with different random initializations to ensure the robustness of the results. MCC = Matthews correlation coefficient; TF = transcription factor; TL = transfer learning.

**Fig. S3**: Performance of MAX models trained with (blue boxes) and without (red boxes) transfer learning on 5,000 (**A**), 1,000 (**B**), 500 (**C**), 250 (**D**), 125 (**E**), and 50 (**F**), bound and unbound regions. Each model was trained five times with different random initializations to ensure the robustness of the results. MCC = Matthews correlation coefficient; TF = transcription factor; TL = transfer learning.

**Fig. S4**: Performance of SPI1 models trained with (blue boxes) and without (red boxes) transfer learning on 5,000 (**A**), 1,000 (**B**), 500 (**C**), 250 (**D**), 125 (**E**), and 50 (**F**), bound and unbound regions. Each model was trained five times with different random initializations to ensure the robustness of the results. MCC = Matthews correlation coefficient; TF = transcription factor; TL = transfer learning.

**Fig. S5**: Performance of SP1 models trained with (blue boxes) and without (red boxes) transfer learning on 5,000 (**A**), 1,000 (**B**), 500 (**C**), 250 (**D**), 125 (**E**), and 50 (**F**), bound and unbound regions. Each model was trained five times with different random initializations to ensure the robustness of the results. MCC = Matthews correlation coefficient; TF = transcription factor; TL = transfer learning.

**Fig S6**: Transfer learning performance using the model architecture of DanQ [46] for the target TFs HNF4A (**A**), JUND (**B**), MAX (**C**), SPI1 (**D**), and SP1 (**E**), from multi-models pre-trained with five TFs with the same binding mode as the target TF (dark blue boxes), five cofactors of the target TF with a different binding mode than the target TF (green boxes), five non-cofactors with the same binding mode as the target TF (light blue boxes), five functional partners of the target TF from STRING with a different binding mode than the target TF (yellow boxes), and five randomly selected TFs with a different binding mode than the target TF (red boxes), with (left) and without (right) the presence of the target TF in the pre-training step. BM = binding mode; MCC = Matthews correlation coefficient; TF = transcription factor.

**Fig. S7**: Transfer learning performance for three target TFs with the same binding mode, but different number of bound regions, from each of the following five families: (**A**) HNF4A, NR2C2, and VD2R (nuclear receptors); (**B**) JUND, ATF7, and NFE2L1 (basic leucine zippers); (**C**) MAX, MNT, and SREBF2 (basic helix-loop-helix factors); (**D**) SPI1, ETV4, and ERG (tryptophan cluster factors); and (**E**) SP1, KLF9, and ZNF740 (C2H2 zinc fingers). Transfer learning performance for each target TF from multi-models pre-trained with five cofactors with different binding modes than the target TF (green boxes), and five randomly selected TFs with a different binding mode than the target TF (red boxes), with (left) and without (right) the presence of the target TF in the pre-training step. The training dataset size of each multi-model is indicated with diamonds (secondary y-axis). The number of bound regions for each TF is shown between parenthesis. BM = binding mode; MCC = Matthews correlation coefficient; TF = transcription factor.

**Fig. S8**: Comparison of Matthews correlation coefficient (MCC; x-axis) with AUCROC (**A**) and AUCPR (**B**) metrics (y-axis) for 148 individual models trained with transfer learning. An MCC >= 0.5, which we define as effective model performance (black dot), corresponds to AUCROC and AUCPR values of ∼0.835.

## Tables

**Table S1:** Total number of ones, zeros and nulls in the sparse matrix for the 163 TFs used in this study.

**Table S2:** Performance for 148 TFs on the original multi-model of 50 TFs, as well as on individual models trained with and without transfer learning. An empty value for multi-model performance indicates that the TF was not used to train the multi-model.

**Table S3:** List of TF binding modes used in this study.

**Table S4:** Tomtom similarities of TF profiles from the JASPAR database to the first convolutional layer filters of the original multi-model of 50 TFs, the individual models of HNF4A and SP1, trained with and without transfer learning, and the individual models of JUND trained with transfer learning from multi-models pre-trained on either five cofactors or five random TFs as well as trained from scratch.

**Table S5:** Transfer learning performance for 148 TFs with and without freezing of the convolutional layers.

## References

1. Lovering RC, Gaudet P, Acencio ML, Ignatchenko A, Jolma A, Fornes O, et al. A GO catalogue of human DNA-binding transcription factors. bioRxiv. Cold Spring Harbor Laboratory; 2020;2020.10.28.359232.

2. Mathelier A, Shi W, Wasserman WW. Identification of altered cis-regulatory elements in human disease. Trends Genet. Elsevier; 2015;31:67–76.

3. Lee R van der, Correard S, Wasserman WW. Deregulated Regulators: Disease-Causing cis Variants in Transcription Factor Genes. Trends Genet. Elsevier; 2020;36:523–39.

4. Nebert DW. Transcription factors and cancer: an overview. Toxicology. 2002;181–182:131–41.

5. Khurana E, Fu Y, Chakravarty D, Demichelis F, Rubin MA, Gerstein M. Role of non-coding sequence variants in cancer. Nat Rev Genet. Nature Publishing Group; 2016;17:93–108.

6. Johnson DS, Mortazavi A, Myers RM, Wold B. Genome-Wide Mapping of in Vivo Protein-DNA Interactions. Science. American Association for the Advancement of Science; 2007;316:1497–502.

7. Chèneby J, Gheorghe M, Artufel M, Mathelier A, Ballester B. ReMap 2018: an updated atlas of regulatory regions from an integrative analysis of DNA-binding ChIP-seq experiments. Nucleic Acids Res. Oxford Academic; 2018;46:D267–75.

8. Chèneby J, Ménétrier Z, Mestdagh M, Rosnet T, Douida A, Rhalloussi W, et al. ReMap 2020: a database of regulatory regions from an integrative analysis of Human and Arabidopsis DNA-binding sequencing experiments. Nucleic Acids Res. Oxford Academic; 2020;48:D180–8.

9. Wasserman WW, Sandelin A. Applied bioinformatics for the identification of regulatory elements. Nat Rev Genet. Nature Publishing Group; 2004;5:276–87.

10. Gheorghe M, Sandve GK, Khan A, Chèneby J, Ballester B, Mathelier A. A map of direct TF–DNA interactions in the human genome. Nucleic Acids Res. Oxford Academic; 2019;47:e21–e21.

11. Snyder MP, Gingeras TR, Moore JE, Weng Z, Gerstein MB, Ren B, et al. Perspectives on ENCODE. Nature. Nature Publishing Group; 2020;583:693–8.

12. Koo PK, Ploenzke M. Deep learning for inferring transcription factor binding sites. Curr Opin Syst Biol [Internet]. 2020 [cited 2020 Jul 10]; Available from: http://www.sciencedirect.com/science/article/pii/S2452310020300032

13. Weiss K, Khoshgoftaar TM, Wang D. A survey of transfer learning. J Big Data. 2016;3:9.

14. Pierson E, Consortium the Gte, Koller D, Battle A, Mostafavi S. Sharing and Specificity of Co-expression Networks across 35 Human Tissues. PLOS Comput Biol. Public Library of Science; 2015;11:e1004220.

15. Yang Y, Fang Q, Shen H-B. Predicting gene regulatory interactions based on spatial gene expression data and deep learning. PLOS Comput Biol. Public Library of Science; 2019;15:e1007324.

16. Mignone P, Pio G, D’Elia D, Ceci M. Exploiting transfer learning for the reconstruction of the human gene regulatory network. Bioinformatics. Oxford Academic; 2020;36:1553–61.

17. Mieth B, Hockley JRF, Görnitz N, Vidovic MM-C, Mü ller K-R, Gutteridge A, et al. Using transfer learning from prior reference knowledge to improve the clustering of single-cell RNA-Seq data. Sci Rep. Nature Publishing Group; 2019;9:20353.

18. Wang J, Agarwal D, Huang M, Hu G, Zhou Z, Ye C, et al. Data denoising with transfer learning in single-cell transcriptomics. Nat Methods. Nature Publishing Group; 2019;16:875–8.

19. Wang T, Johnson TS, Shao W, Lu Z, Helm BR, Zhang J, et al. BERMUDA: a novel deep transfer learning method for single-cell RNA sequencing batch correction reveals hidden high-resolution cellular subtypes. Genome Biol. 2019;20:165.

20. Lotfollahi M, Naghipourfar M, Luecken MD, Khajavi M, Büttner M, Avsec Z, et al. Query to reference single-cell integration with transfer learning. bioRxiv. Cold Spring Harbor Laboratory; 2020;2020.07.16.205997.

21. Avsec Ž, Kreuzhuber R, Israeli J, Xu N, Cheng J, Shrikumar A, et al. The Kipoi repository accelerates community exchange and reuse of predictive models for genomics. Nat Biotechnol. Nature Publishing Group; 2019;37:592–600.

22. Schwessinger R, Gosden M, Downes D, Brown RC, Oudelaar AM, Telenius J, et al. DeepC: predicting 3D genome folding using megabase-scale transfer learning. Nat Methods. Nature Publishing Group; 2020;17:1118–24.

23. Lan G, Zhou J, Xu R, Lu Q, Wang H. Cross-Cell-Type Prediction of TF-Binding Site by Integrating Convolutional Neural Network and Adversarial Network. Int J Mol Sci. Multidisciplinary Digital Publishing Institute; 2019;20:3425.

24. Zheng A, Lamkin M, Wu C, Su H, Gymrek M. Deep neural networks identify context-specific determinants of transcription factor binding affinity. bioRxiv. Cold Spring Harbor Laboratory; 2020;2020.02.26.965343.

25. Deng J, Dong W, Socher R, Li L, Kai Li, Li Fei-Fei. ImageNet: A large-scale hierarchical image database. 2009 IEEE Conf Comput Vis Pattern Recognit. 2009. p. 248–55.

26. Zeiler MD, Fergus R. Visualizing and Understanding Convolutional Networks. ArXiv13112901 Cs [Internet]. 2013 [cited 2020 Oct 27]; Available from: http://arxiv.org/abs/1311.2901

27. Nakato R, Shirahige K. Recent advances in ChIP-seq analysis: from quality management to whole-genome annotation. Brief Bioinform. Oxford Academic; 2017;18:279–90.

28. Karimzadeh M, Hoffman MM. Virtual ChIP-seq: predicting transcription factor binding by learning from the transcriptome. bioRxiv. Cold Spring Harbor Laboratory; 2019;168419.

29. Bailey TL, Machanick P. Inferring direct DNA binding from ChIP-seq. Nucleic Acids Res. 2012;40:e128.

30. Wang J, Zhuang J, Iyer S, Lin X, Whitfield TW, Greven MC, et al. Sequence features and chromatin structure around the genomic regions bound by 119 human transcription factors. Genome Res. 2012;22:1798–812.

31. Teytelman L, Thurtle DM, Rine J, Oudenaarden A van. Highly expressed loci are vulnerable to misleading ChIP localization of multiple unrelated proteins. Proc Natl Acad Sci. National Academy of Sciences; 2013;110:18602–7.

32. Worsley Hunt R, Wasserman WW. Non-targeted transcription factors motifs are a systemic component of ChIP-seq datasets. Genome Biol [Internet]. 2014 [cited 2020 Jul 21];15. Available from: https://www.ncbi.nlm.nih.gov/pmc/articles/PMC4165360/

33. Wreczycka K, Franke V, Uyar B, Wurmus R, Bulut S, Tursun B, et al. HOT or not: examining the basis of high-occupancy target regions. Nucleic Acids Res. Oxford Academic; 2019;47:5735–45.

34. Dror I, Golan T, Levy C, Rohs R, Mandel-Gutfreund Y. A widespread role of the motif environment in transcription factor binding across diverse protein families. Genome Res. 2015;25:1268–80.

35. Worsley Hunt R, Mathelier A, del Peso L, Wasserman WW. Improving analysis of transcription factor binding sites within ChIP-Seq data based on topological motif enrichment. BMC Genomics. 2014;15:472.

36. Frenkel ZM, Trifonov EN, Volkovich Z, Bettecken T. Nucleosome Positioning Patterns Derived from Human Apoptotic Nucleosomes. J Biomol Struct Dyn. Taylor & Francis; 2011;29:577–83.

37. Zhu F, Farnung L, Kaasinen E, Sahu B, Yin Y, Wei B, et al. The interaction landscape between transcription factors and the nucleosome. Nature. Nature Publishing Group; 2018;562:76–81.

38. Eraslan G, Avsec Ž, Gagneur J, Theis FJ. Deep learning: new computational modelling techniques for genomics. Nat Rev Genet. Nature Publishing Group; 2019;20:389–403.

39. Wingender E, Schoeps T, Haubrock M, Krull M, Dönitz J. TFClass: expanding the classification of human transcription factors to their mammalian orthologs. Nucleic Acids Res. Oxford Academic; 2018;46:D343–7.

40. Capellera-Garcia S, Pulecio J, Dhulipala K, Siva K, Rayon-Estrada V, Singbrant S, et al. Defining the Minimal Factors Required for Erythropoiesis through Direct Lineage Conversion. Cell Rep. Elsevier; 2016;15:2550–62.

41. Lu R, Mucaki EJ, Rogan PK. Discovery and validation of information theory-based transcription factor and cofactor binding site motifs. Nucleic Acids Res. Oxford Academic; 2017;45:e27–e27.

42. Szklarczyk D, Gable AL, Lyon D, Junge A, Wyder S, Huerta-Cepas J, et al. STRING v11: protein–protein association networks with increased coverage, supporting functional discovery in genome-wide experimental datasets. Nucleic Acids Res. Oxford Academic; 2019;47:D607–13.

43. Chicco D, Jurman G. The advantages of the Matthews correlation coefficient (MCC) over F1 score and accuracy in binary classification evaluation. BMC Genomics. 2020;21:6.

44. Ambrosini G, Vorontsov I, Penzar D, Groux R, Fornes O, Nikolaeva DD, et al. Insights gained from a comprehensive all-against-all transcription factor binding motif benchmarking study. Genome Biol. 2020;21:114.

45. Fornes O, Castro-Mondragon JA, Khan A, van der Lee R, Zhang X, Richmond PA, et al. JASPAR 2020: update of the open-access database of transcription factor binding profiles. Nucleic Acids Res. Oxford Academic; 2020;48:D87–92.

46. Quang D, Xie X. DanQ: a hybrid convolutional and recurrent deep neural network for quantifying the function of DNA sequences. Nucleic Acids Res. Oxford Academic; 2016;44:e107–e107.

47. Gupta S, Stamatoyannopoulos JA, Bailey TL, Noble WS. Quantifying similarity between motifs. Genome Biol. 2007;8:R24.

48. Shrikumar A, Greenside P, Kundaje A. Learning Important Features Through Propagating Activation Differences. ArXiv170402685 Cs [Internet]. 2019 [cited 2020 Oct 26]; Available from: http://arxiv.org/abs/1704.02685

49. Shrikumar A, Tian K, Avsec Ž, Shcherbina A, Banerjee A, Sharmin M, et al. Technical Note on Transcription Factor Motif Discovery from Importance Scores (TF-MoDISco) version 0.5.6.5. ArXiv181100416 Cs Q-Bio Stat [Internet]. 2020 [cited 2020 Oct 26]; Available from: http://arxiv.org/abs/1811.00416

50. Sandelin A, Wasserman WW. Constrained binding site diversity within families of transcription factors enhances pattern discovery bioinformatics. J Mol Biol. 2004;338:207–15.

51. Smith SL, Kindermans P-J, Ying C, Le Qv. Don’t Decay the Learning Rate, Increase the Batch Size. ArXiv171100489 Cs Stat [Internet]. 2018 [cited 2020 Dec 18]; Available from: http://arxiv.org/abs/1711.00489

52. Kandel I, Castelli M. The effect of batch size on the generalizability of the convolutional neural networks on a histopathology dataset. ICT Express. 2020;6:312–5.

53. Dunham I, Kundaje A, Aldred SF, Collins PJ, Davis CA, Doyle F, et al. An integrated encyclopedia of DNA elements in the human genome. Nature. Nature Publishing Group; 2012;489:57–74.

54. Lee CM, Barber GP, Casper J, Clawson H, Diekhans M, Gonzalez JN, et al. UCSC Genome Browser enters 20th year. Nucleic Acids Res. Oxford Academic; 2020;48:D756–61.

55. Quinlan AR, Hall IM. BEDTools: a flexible suite of utilities for comparing genomic features. Bioinformatics. Oxford Academic; 2010;26:841–2.

56. Pedregosa F, Varoquaux G, Gramfort A, Michel V, Thirion B, Grisel O, et al. Scikit-learn: Machine Learning in Python. J Mach Learn Res. 2011;12:2825–30.

57. Kelley DR, Snoek J, Rinn JL. Basset: learning the regulatory code of the accessible genome with deep convolutional neural networks. Genome Res. 2016;26:990–9.

58. Maslova A, Ramirez RN, Ma K, Schmutz H, Wang C, Fox C, et al. Deep learning of immune cell differentiation. Proc Natl Acad Sci. National Academy of Sciences; 2020;117:25655–66.

59. Paszke A, Gross S, Massa F, Lerer A, Bradbury J, Chanan G, et al. PyTorch: An Imperative Style, High-Performance Deep Learning Library. Adv Neural Inf Process Syst. 2019;32:8026–37.

60. Kingma DP, Ba J. Adam: A Method for Stochastic Optimization. ArXiv14126980 Cs [Internet]. 2017 [cited 2020 Jul 10]; Available from: http://arxiv.org/abs/1412.6980

61. Zhou J, Troyanskaya OG. Predicting effects of noncoding variants with deep learning–based sequence model. Nat Methods. Nature Publishing Group; 2015;12:931–4.

62. Kokhlikyan N, Miglani V, Martin M, Wang E, Alsallakh B, Reynolds J, et al. Captum: A unified and generic model interpretability library for PyTorch. ArXiv200907896 Cs Stat [Internet]. 2020 [cited 2020 Nov 12]; Available from: http://arxiv.org/abs/2009.07896

63. Harris CR, Millman KJ, van der Walt SJ, Gommers R, Virtanen P, Cournapeau D, et al. Array programming with NumPy. Nature. Nature Publishing Group; 2020;585:357–62.

