## Supplementary figures and images for "Biologically-relevant transfer learning improves transcription factor binding prediction"

### Fig.S1.png

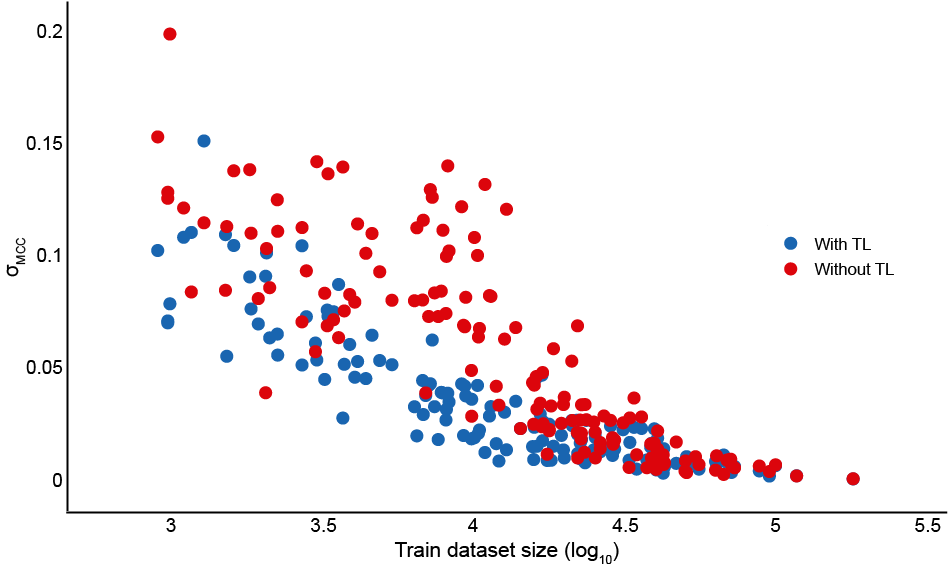

### Fig.S2.png

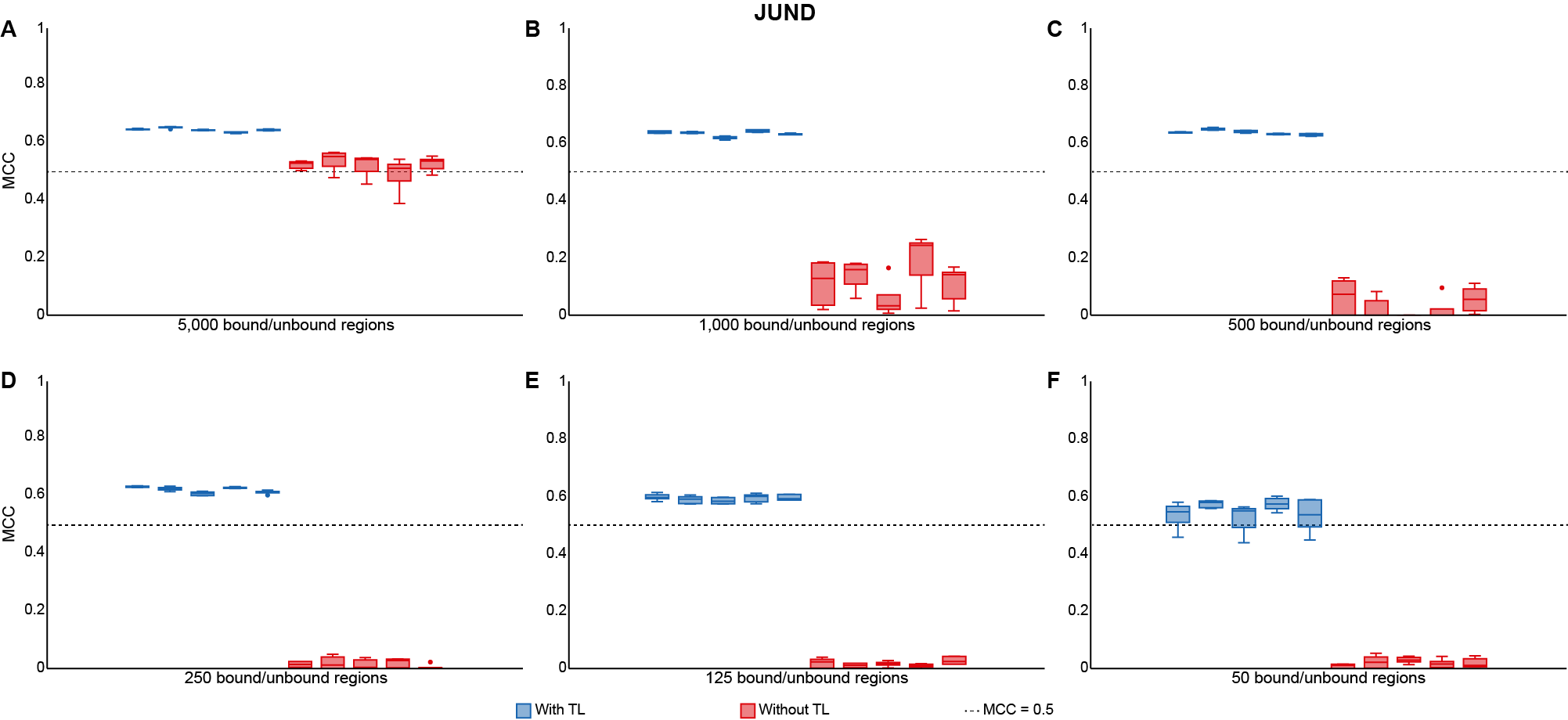

### Fig.S3.png

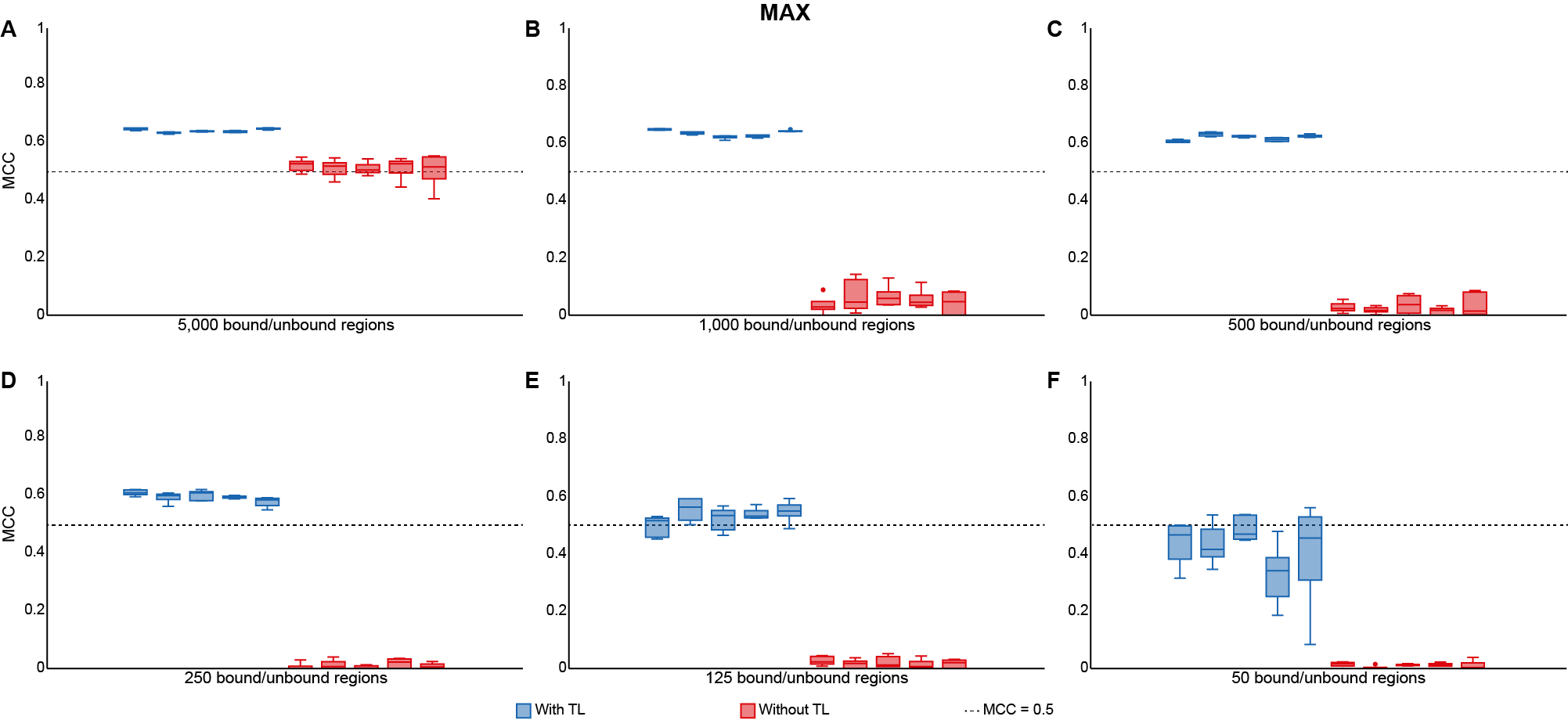

### Fig.S4.png

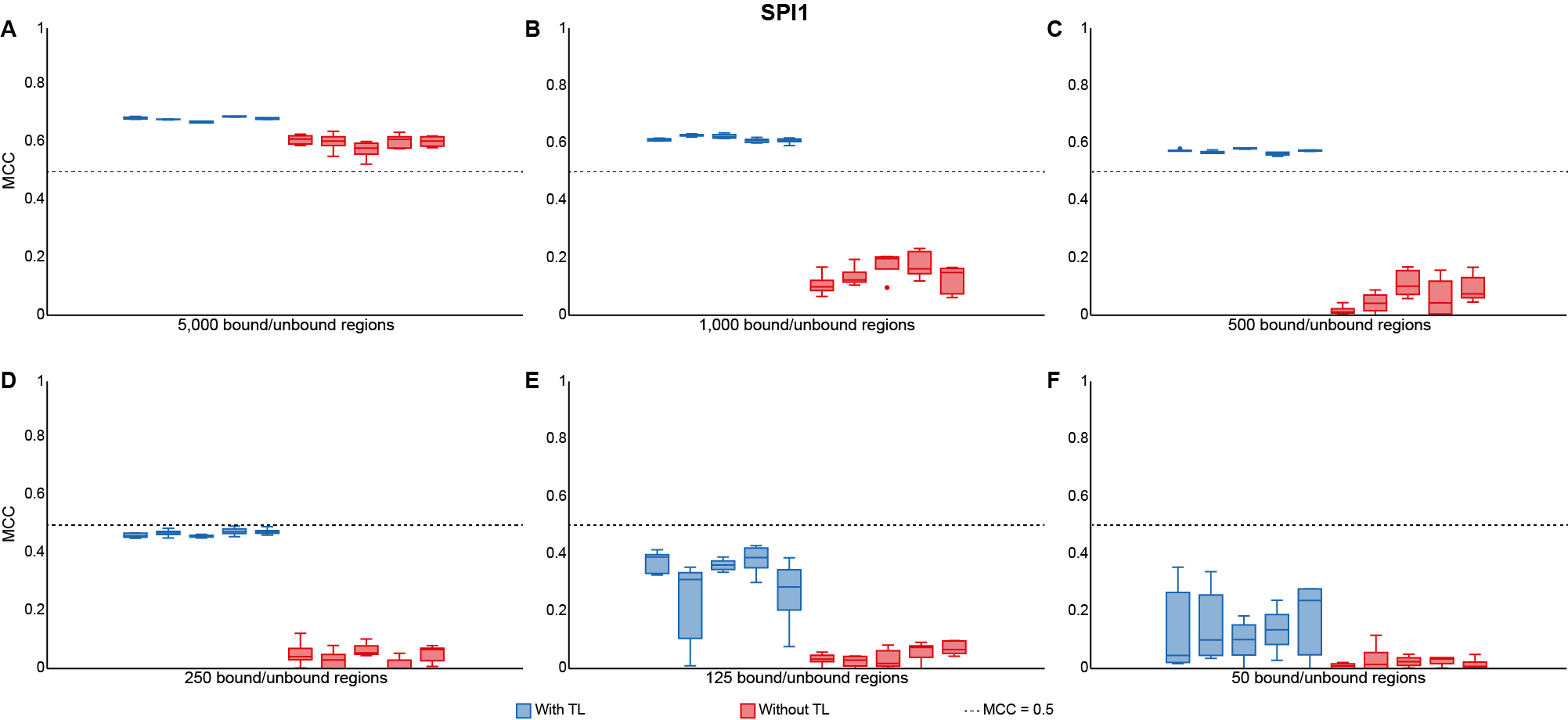

### Fig.S5.png

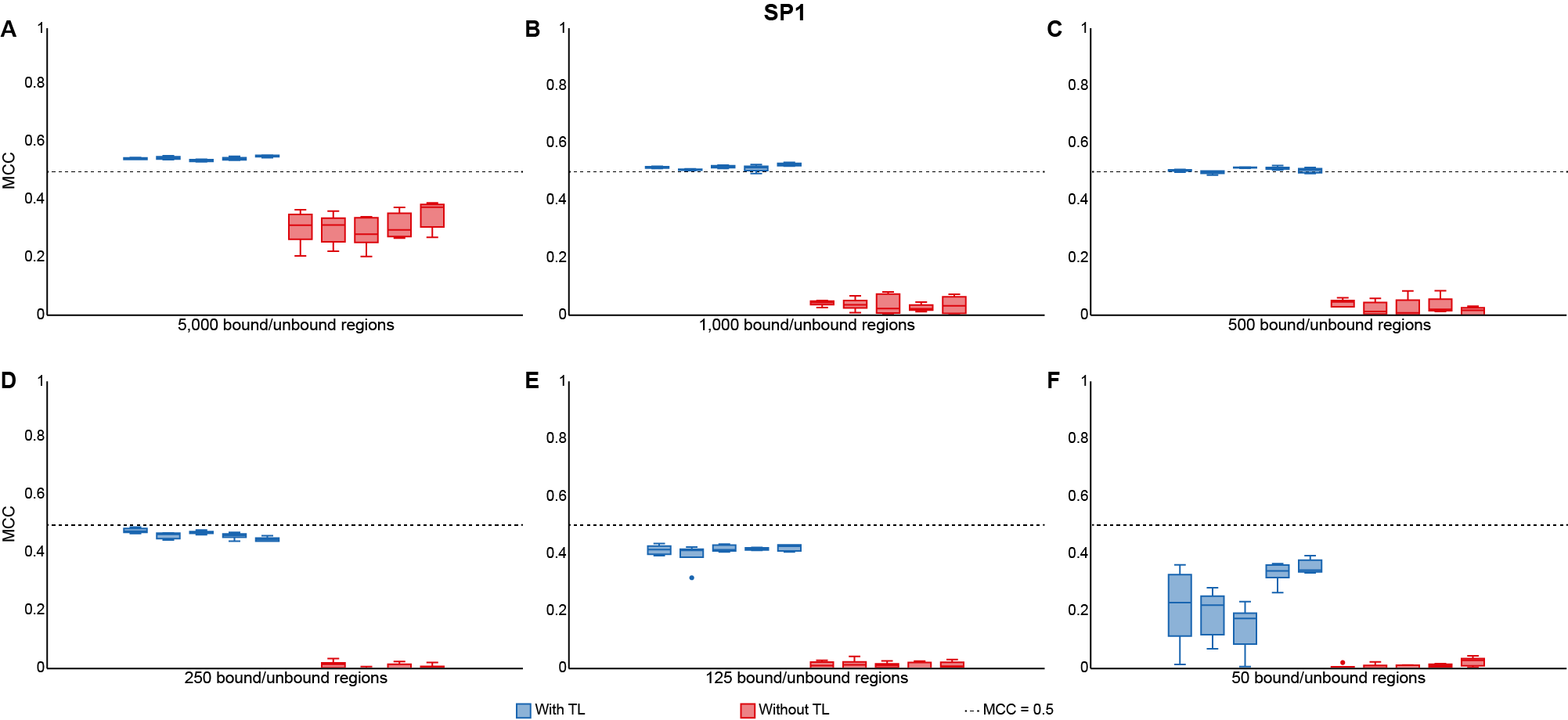

### Fig.S6.png

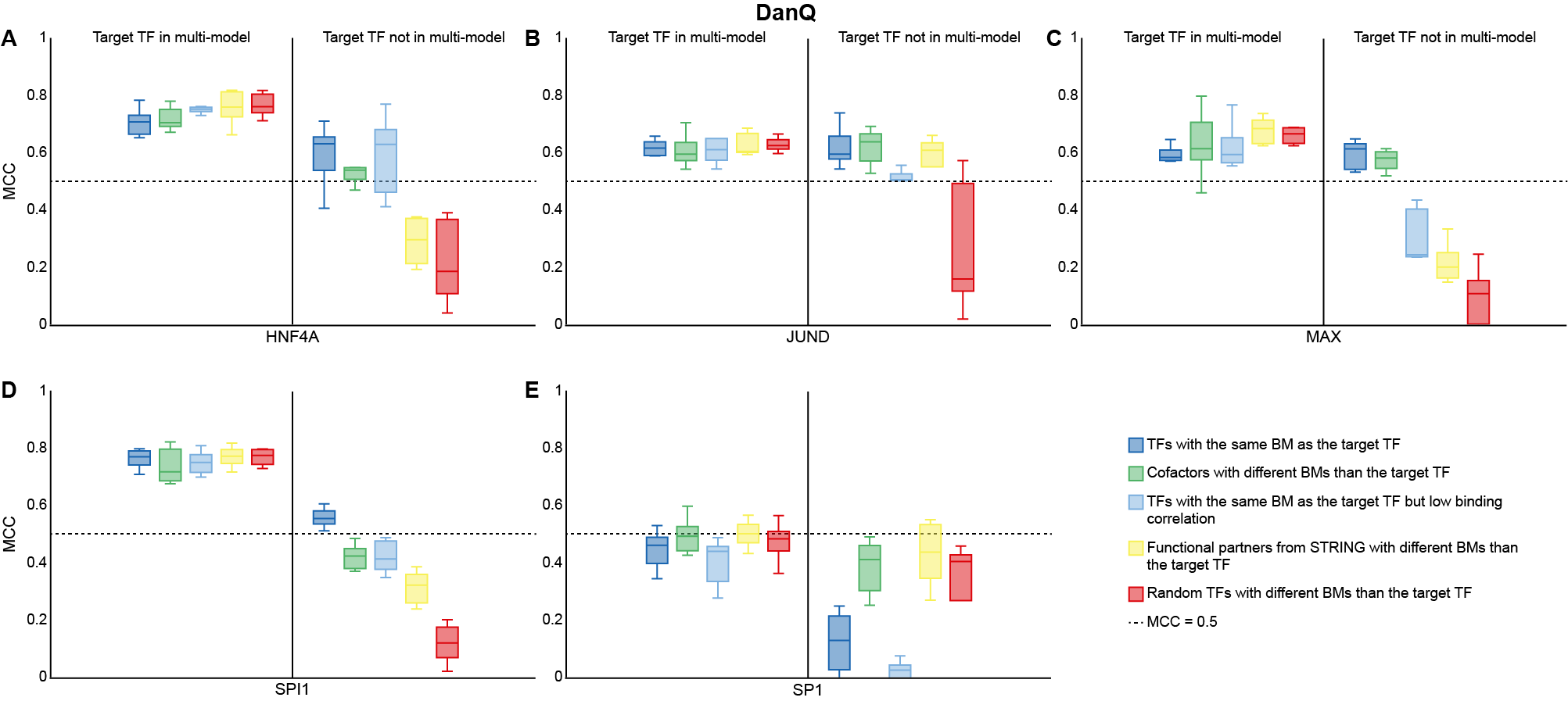

### Fig.S7.png

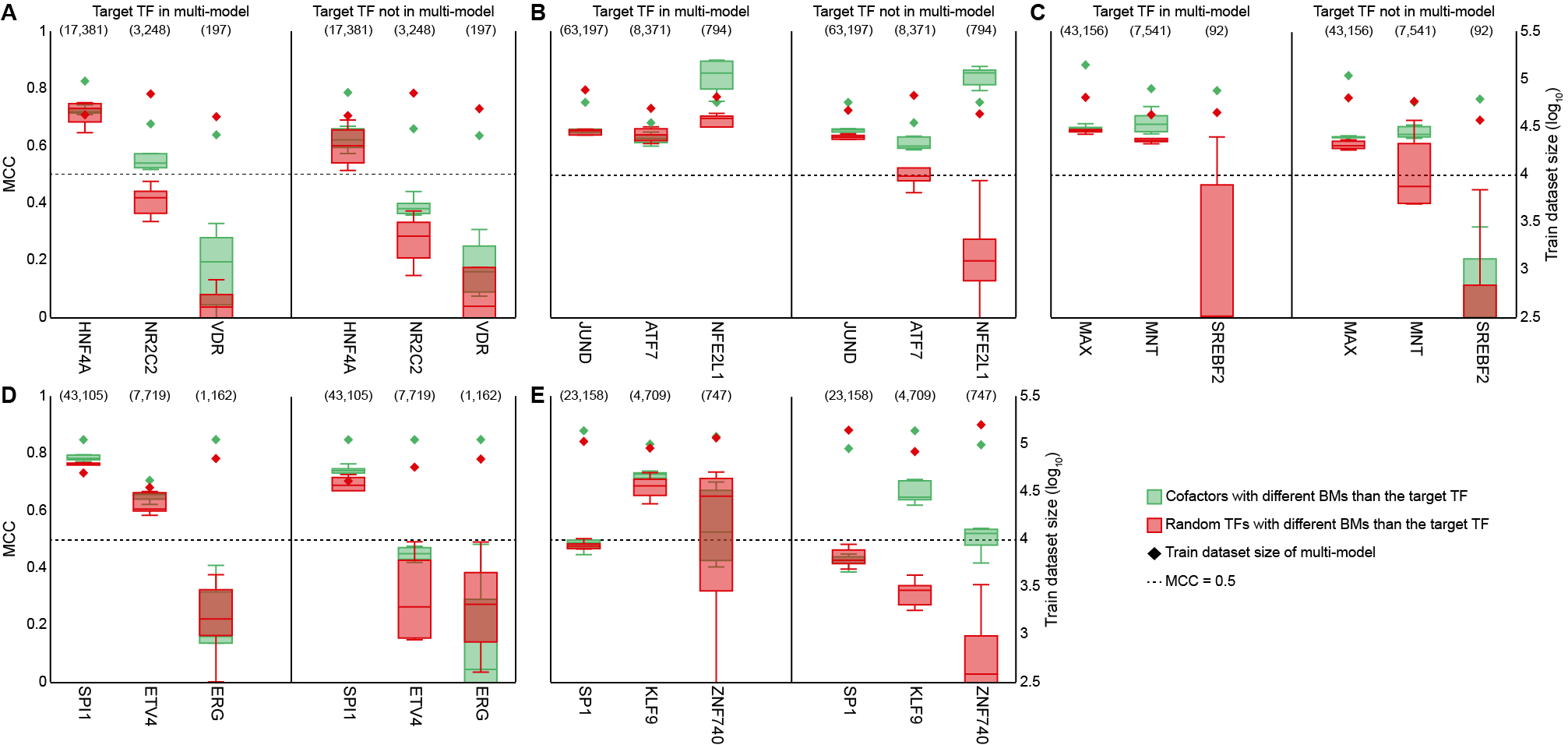

### Fig.S8.png

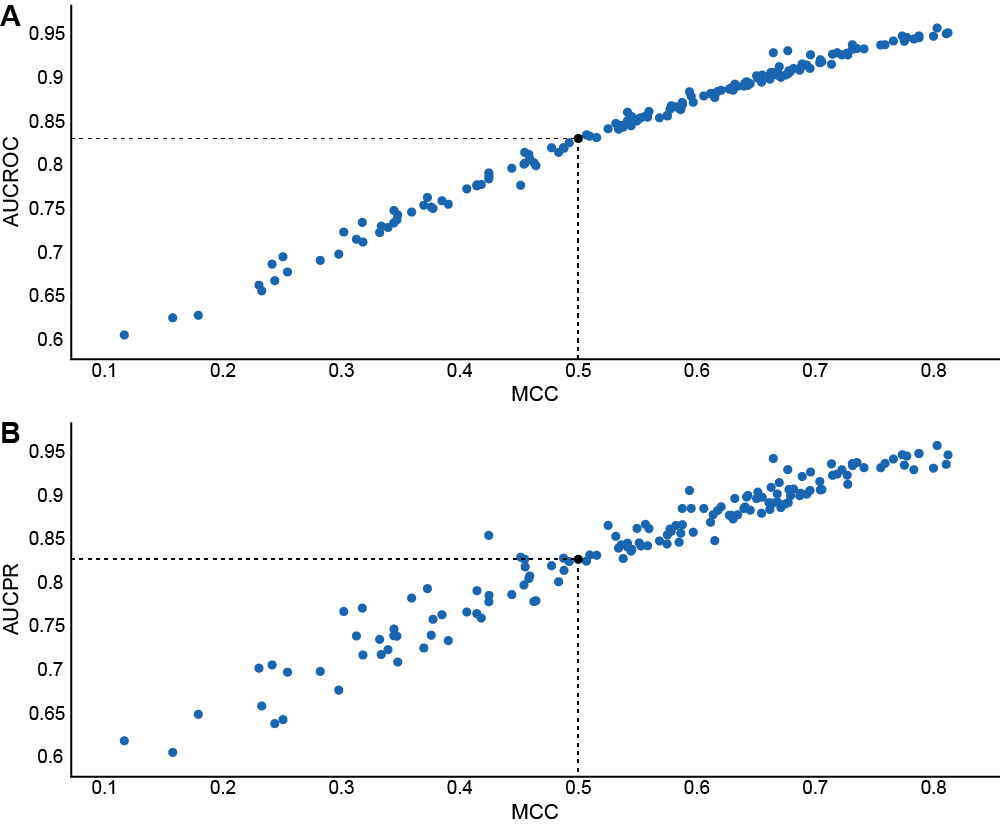
